# Paused Polymerase synergises with transcription factors and RNA to enhance the transcriptional potential of promoters

**DOI:** 10.1101/2024.10.18.618990

**Authors:** Rajat Mann, Deepanshu Soota, Ayan Das, Nitesh Kumar Podh, Gunjan Mehta, Dimple Notani

## Abstract

Regulated pausing of RNA polymerase II (Pol II) is essential for enabling rapid and coordinated transcriptional responses to signalling cues. Pausing also contributes to the formation of nucleosome-free regions with the help of chromatin remodellers. However, if these nucleosome-free regions engage with transcription factors to stimulate the transcription potential of paused promoters is not known. In this study, we demonstrate that ligand-induced estrogen receptor-alpha (ERα) binding is stabilized at Pol II-paused sites. This stabilization results from an increased dwell time of ERα on chromatin, as revealed by single molecule tracking (SMT) experiments. Notably, short chromatin-associated RNAs generated by the paused Pol II contribute to enhancing ERα binding at paused promoters. We also observe that pausing increases histone H3K27 acetylation (H3K27ac) levels, which primes paused promoters for robust transcriptional activation upon release. Collectively, these findings suggest that paused Pol II plays a central role in enhancing transcription factor binding through an RNA-dependent mechanism. This, in turn, results in a more vigorous transcriptional response following pausing release, thus contributing to the fine-tuning of ERα-mediated gene regulation.

## Introduction

Transcription factors (TFs) regulate gene expression by binding to their cognate DNA motifs. Upon binding, they recruit cofactors and RNA polymerases to the site for transcription. Once the polymerase is recruited and the pre-initiation complex (PIC) is formed, the polymerase transcribes RNA (Aibara et al., 2021). In around 30-50% of genes, the polymerase remains paused near the TSS after synthesizing around 20-60 bases of RNA (Core et al., 2008; Muse et al., 2007; Zeitlinger et al., 2007). This promoter-proximal pausing is one of the major rate-limiting steps in transcription (Gressel et al., 2019; Liu et al., 2014). The release of the paused polymerase into productive elongation is a highly regulated process involving multiple players like DNA sequence, NELF, pTEFb, Spt5, etc. (Adelman & Lis, 2012; Core & Adelman, 2019; Vos et al., 2018).

In addition, the polymerase can also recruit chromatin-remodellers to the paused sites resulting in creation of unstable nucleosomes that are dynamic (Brahma & Henikoff, 2024; Materne et al., 2015; Soutourina et al., 2006; Yague-Sanz et al., 2023). Recent studies have highlighted the role of BAF complex, in destabilizing the nucleosome at the polymerase paused sites. The transcription factor binding at exposed TF motif around these unstable nucleosomes can further recruit these remodelers complexes leading to stable NFR formation (Brahma & Henikoff, 2024). Besides creating open permissive chromatin, the paused polymerase also transcribes short RNA from the TSS (Henriques et al., 2013; Nechaev et al., 2010). The short RNA that is stably associated with polymerase is protected from exonuclease digestion, while the ones released get degraded fast (Henriques et al., 2013). Therefore, Nucleosome-free regions (NFR) and short RNA are the primary products at paused polymerase sites (Gilchrist et al., 2008, 2010; Henriques et al., 2013, 2013; Nechaev et al., 2010). However, it is not known if there is a functional link between transcription factor binding, short RNA and transcriptional potential of paused genes after release.

A vast majority of transcription factors are known to interact with RNA through RNA binding domains (Mann & Notani, 2023; Oksuz et al., 2023; Soota et al., 2024). The binding of TF with RNA can significantly alter their binding affinity for cognate DNA motifs (Oksuz et al., 2023; Soota et al., 2024). We speculated that the NFRs created due to the action of paused polymerase and remodellers may expose additional TF binding sites, facilitating TF recruitment. In addition, the short RNA produced from the paused polymerase can add to this by stabilizing TF binding. Further, this increased TF recruitment, along with the regulatory machinery of interacting TFs and co-activators that it brings, may increase the transcriptional activation of the gene upon release from inhibition.

Towards this, we utilized estrogen signalling induced transcription factor ERα, which is also known to bind to RNA (Oksuz et al., 2023; Soota et al., 2024; Xu et al., 2021). We reveal that the binding of ERα on regulatory regions increases upon chemical-induced transcription elongation inhibition (DRB: 5,6-Dichloro-1-β-D-ribofuranosylbenzimidazole). This increase was correlated with enhanced dwell time of ERα detected by single molecule tracking (SMT) and resulting in the formation of bigger ERα condensates. Moreover, the increased binding was dependent on the short RNA from the paused polymerase as treating cells with triptolide, which degrades the polymerase, led to reduced ERα binding. Whereas chromatin-associated RNAs physically interacted with ERα, stabilizing its binding at paused sites. As a consequence pausing, the nucleosomes were hyperacetylated, thereby increasing the rate of gene transcription upon release of pausing. These results reveal that paused promoters enhance their transcription potential by virtue of interaction between transcription factors and short RNAs.

## Results

### Transcription elongation inhibition enhances ERα binding on chromatin

The response to stimulation, such as hormone signalling is prompt (Hah et al., 2011; Falo-Sanjuan et al., 2019; Keenan et al., 2020). The fast transcriptional response is regulated by the release of polymerase complex from pausing (Bartman et al., 2019). In order to understand the role of polymerase pausing on transcription factor binding, we first treated MCF-7 breast cancer cells with transcription elongation inhibitor DRB. DRB targets the CDK9 subunit of the pTEFb complex which is important in pause release and leads to its inhibition. Then, we stimulated the binding of ERα in the genome by exposing the cell to oestradiol, E2 for one hour (Fig 1A). We first checked the effectiveness of treatment by performing qPCR and observed that both myc and pre-myc were significantly down-regulated upon DRB, but ERα levels were unaffected (Fig S1A) treatment. We then performed ChIP-seq for ERα following the same treatment regime. A significant increase in the ERα binding was observed in the genome (Fig. 1B-C). Approximately 2271 new binding events occurred, and 1493 sites were lost. Though, 6574 peaks remained constant but they also showed increased binding of ERα upon DRB treatment (Fig 1C-D).

**Figure 1:**
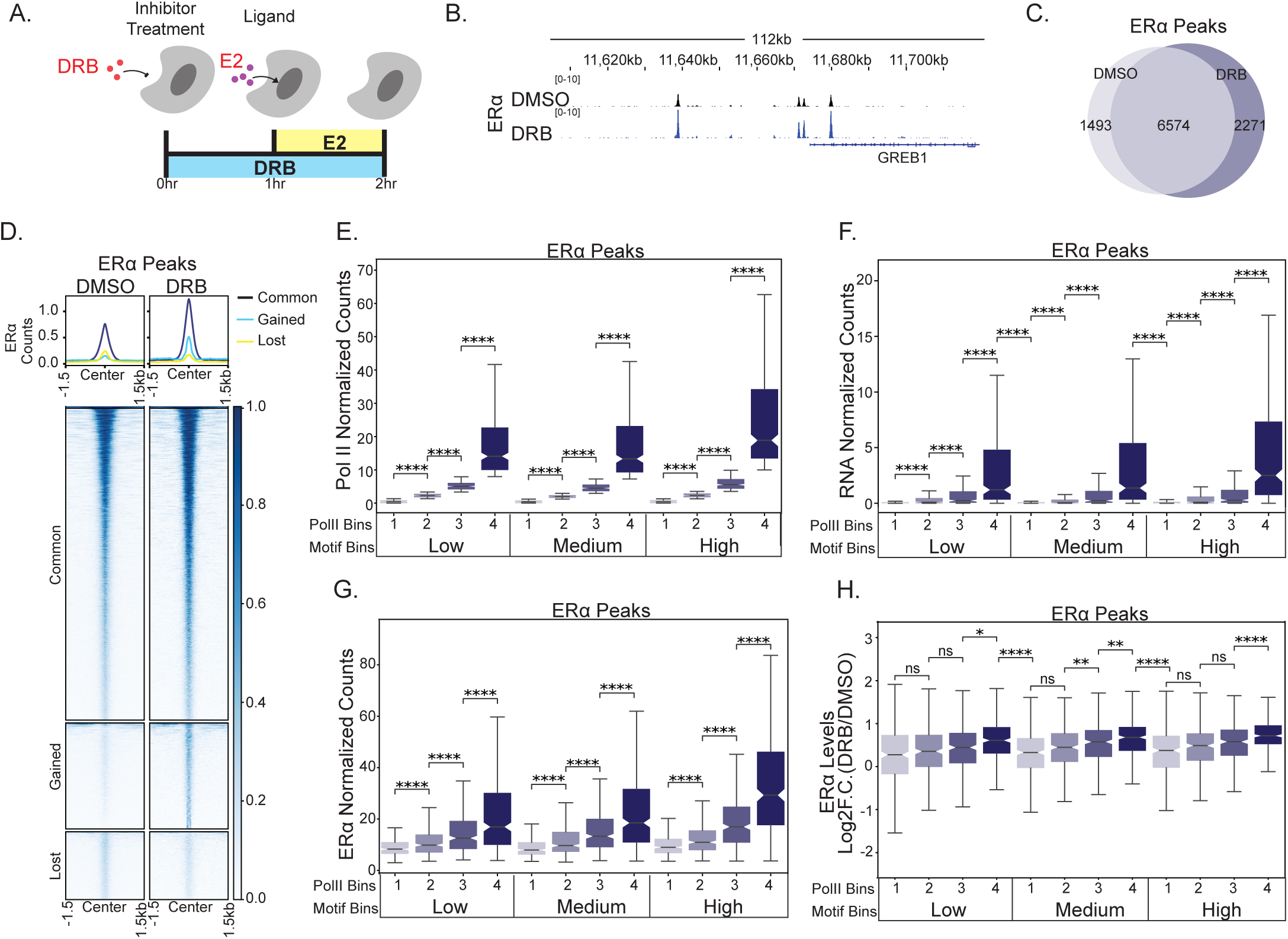
Transcription elongation inhibition enhances ERα binding on chromatin. **A.** Schematic showing the treatment regime for transcription inhibition and E2 treatment. **B.** IgV snapshot of ERα binding on GREB1 gene locus in DMSO and DRB. **C.** Venn diagram showing loss/gain/common ERα peaks upon DRB treatment. D. Heatmap showing the binding of ERα in common, gained and lost sites upon DRB treatment. E, F and G Pol II, RNA, and ERα levels respectively on the ERα peaks sorted first based on the motif scores into low, medium and high. Followed by dividing each category into four bins based on the Pol II levels from low (1)-to high (4). p-value is calculated using the Mann-Whitney-Wilcoxon test. **H.** Boxplot showing a gain in the binding of ERα (log2 fold change (DRB/DMSO) on peaks in the categories sorted based on motif and Pol II levels. p-value is calculated using the Mann-Whitney-Wilcoxon test.

To understand the effect of polymerase levels on top of the motif strength in governing the increased ERα binding upon transcription inhibition, we first divided the ERα peaks based on their motif score into low, medium and high. Since, motif strength is a primary regulator of TF binding (Dror et al., 2015; Inukai et al., 2017). We segregated each motif bin into four sub-bins in increasing order of Pol II levels from 1(low) to 4(high) and confirmed the transcription levels at these bins (Fig 1E-F). We then, compared the binding of ERα within each motif bin. Even though the motif score for the categories was similar (Fig S1B), the bins showing higher PolII levels showed better binding with ERα (Fig 1G) which is expected. After treating the cells with DRB, the binding of ERα still remained majorly dependent on Pol II levels (Fig S1C). Further, ERα peaks that already exhibited high Pol II were the ones showing the highest gain in ERα binding upon DRB treatment (Fig 1H). The data suggests that paused pol II allows more binding of ERα, and the level of ERα gain depends on the level of pre-existing pol II.

### Transcription elongation inhibition leads to bigger and fast forming ERα condensates

ERα forms phase-separated condensates, which are heterogenous assemblies of ERα, mediator complex and possibly other factors (Nair et al., 2019; Saravanan et al., 2020). We observed that these condensates were formed in an E2-dependent manner and corroborated with the transcriptional response of estrogen signalling. The ERα condensates formation peaked as the transcriptional response to signalling peaks at 60 mins post-E2 addition, and drops as the transcriptional response decreases (120-180 mins) (Fig S2A-B). We then tested the overlap between condensates and RNA Pol II. Towards this, the normalized overlap between ERα and Pol IIS5P spots was calculated and found to be increasing in the same order as ERα condensates during the signalling time course (Fig S2C). Similarly, we also tested the overlap between ERα condensates and nascent transcription spots and observed the spatial overlap between them and they also followed the same pattern as Pol IIS5P, i.e. peaking at 60 mins and then decreasing at 180 mins (Fig S2D-E). The data suggests that ERα condensates correlate with active polymerase as well as nascent transcription.

Next, to understand the effect of polymerase pausing on these Pol II-containing ERα condensates, we performed live-cell imaging of ERα-GFP in MCF-7 cells after DRB treatment following the same regime as shown (Fig 1A) and imaged the cells after adding E2 for 60 mins. Before E2 addition, ERα-GFP was distributed homogeneously inside the nucleus but the ERα condensates emerged within 5-10 minutes after E2 addition (Fig 2A). The size of ERα condensates (quantified using average diameter) and the number (quantified using number density) increased exponentially upon E2 addition but reached a plateau phase around 40 minutes (Fig 2B-C). We quantified the rate of their increase by calculating the t-half and the maxima for each curve (Fig S2F-I). Upon treating the cells with DRB, the maximum number density did not show any significant change, but the t-half for the curve showed a faster rate of increase in the condensate numbers (Fig 2D-E). We also measured the average diameter of condensate upon transcription inhibition and observed a significant increase in the size of condensates even though ERα concentration was similar (Fig 2F, Fig S2J). However, the rate of size increase of ERα condensate was not significantly different between DMSO and DRB (Fig 2G). The data suggests that transcription elongation inhibition allows faster formation of condensates with increased size, which corroborates with the higher ERα binding in the genome, as seen in Fig. 1.

**Figure 2:**
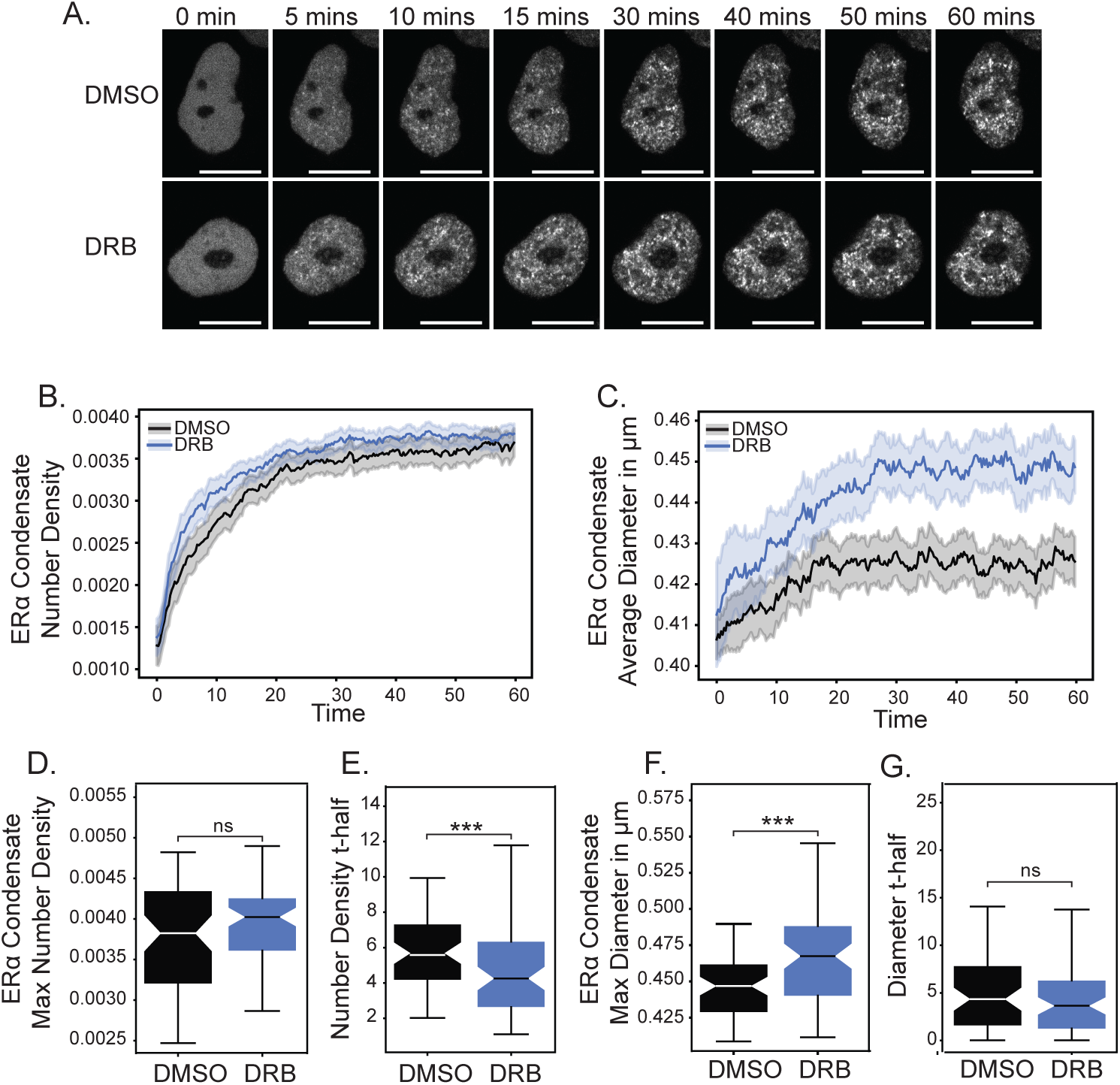
Transcription elongation inhibition leads to bigger and fast forming ERα condensates. **A.** Representative images of MCF-7 cell nucleus tagged with ERα-eGFP imaged from 0 to 60mins. Frame rate: 15s/frame. **B and C.** Plot of number density and average diameter of ERα condensates with time for DMSO and DRB. Solid lines indicate the median curve, and the hue represents a 95% confidence interval. (n=64 and 75 for DMSO and DRB, respectively). **D and F.** Boxplot of max number density and max averaged diameter for ERα condensates in DMSO and DRB conditions. **E and G.** Boxplot of t-half in min calculated for number density and averaged diameter in DMSO and DRB conditions.

### Single molecule tracking revealed higher bound fraction and residence time of ERα on chromatin upon Pol II pausing

As the high ERα binding was observed upon DRB treatment by ChIP-seq analysis, we quantified the fraction of bound molecules of ERα in the nucleus and its residence time using SMT (Single Molecule Tracking). We expressed ERα-HaloTag from the EFS promoter and labelled it using JF646-HaloTag Ligand (HTL) (Fig 3A). Single molecule imaging and tracking was performed as described previously (Podh et al. 2022, Swinstead et al. 2016). Time-lapse movies were acquired with two imaging regimes: 1) fast imaging regime (15 ms time interval for 1000 frames) to quantify the diffusion parameters (fraction of bound and unbound molecules) and 2) slow imaging regime (200 ms time interval for 600 frames) to quantify the residence time. We observed that upon DRB treatment, the bound fraction of ERα increased from 68.4% (DMSO) to 76.7% (DRB, Fig 3B). The residence time of the specific bound fraction (yellow fraction in the pie chart) also increased from 10.3±1.0 s (DMSO) to 13.4±1.4 s (DRB, p<0.05 by z test, Fig 3C). These results suggest that the DRB treatment increases the bound fraction of ERα and its residence time. To further validate that the stronger binding of ERα leads to the formation of bigger chromatin-bound condensate, we treated cells with CSK, which removes the free-floating protein from the nucleus (Gu et al., 2020). Then, immunostained the cells for ERα and calculated the volume fraction for the ERα condensates (Fig 3D). Similar to live cells, we found an increase in the chromatin-bound ERα condensate formation after DRB treatment (Fig 3E). Together, these results suggest that transcription inhibition leads to bigger chromatin-bound ERα condensates as well as an increase in the dwell time of stably bound ERα.

**Figure 3.**
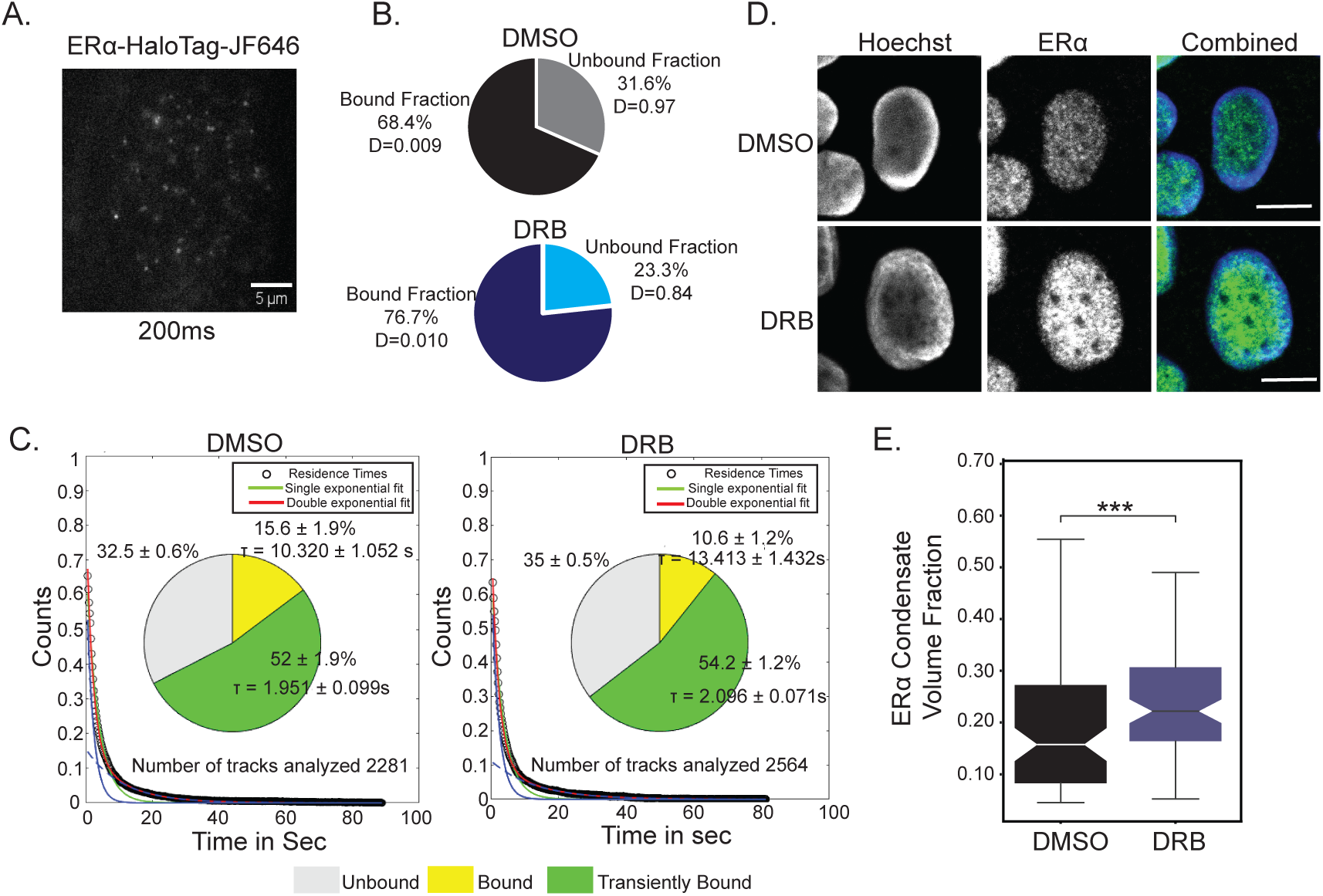
Single molecule tracking revealed higher bound fraction and residence time of ERα on chromatin upon Pol II pausing. **A.** Representative image shows the single molecules of ERα-HaloTag-JF646-HTL in the nucleus of MCF7 cells. **B.** Spot-On based kinetic modelling for the robust quantification of the fraction of bound and unbound molecules, quantified from movies acquired with fast imaging regime (15 ms time-interval). Pie charts represent the fraction of chromatin-bound and unbound molecules derived from modelling CDFs over 15-120 ms intervals and their mean diffusion coefficient. **C.** Survival time distribution of ERα in the presence and absence of DRB quantified from the movies acquired with slow imaging regime (200 ms time-interval). The distributions fit well with the double exponential curve (red line), suggesting two types of bound population: 1) specific bound fraction (with long residence time) and 2) non-specifically/transiently bound fraction (with short residence time). Pie charts represent the percentage of molecules specifically bound (yellow), non-specifically/transiently bound (green) and unbound (grey). The mean residence times (T) of specific and non-specific fractions are presented next to their representative fractions. **D.** Representative images of MCF-7 nucleus after treatment with CSK buffer and immunostained with ERα in DMSO and DRB. **E.** Plot of volume fraction of ERα condensates from CSK-treated images in DMSO and DRB.

### RNA from the pol II paused sites contributes to ERα binding

The paused polymerase transcribes short RNA, and they are present in abundance. In principle, chromatin accessible regions created by paused polymerase and the RNA-TF interaction with the short RNA both could stabilize ERα binding. We then decided to decouple whether the polymerase complex or RNA is responsible for the increased ERα binding upon DRB treatment. We first degraded the polymerase using triptolide (TRP), which also led to a decrease in RNA levels by inhibiting the transcription initiation (Fig 4A). We treated the cells with TRP following the same regime as in the case of DRB treatment and assessed the binding of ERα (Fig 1A). The ChIP-seq of ERα revealed a widespread loss of ERα binding in the genome (Fig 4B, C and S3E). Further, we observed that the sites with the highest Pol II were the most affected by TRP treatment (Fig 4D and S3A). The data suggest that RNA Pol II complex and chromatin RNA are both required for gain in ERα binding. Though, both the TRP and DRB treatments reduce the levels of chromatin-bound RNA. However, DRB treatment does not inhibit transcription initiation. Short RNA accumulates around paused Pol II sites (Henriques et al., 2013; Nechaev et al., 2010).

**Figure 4:**
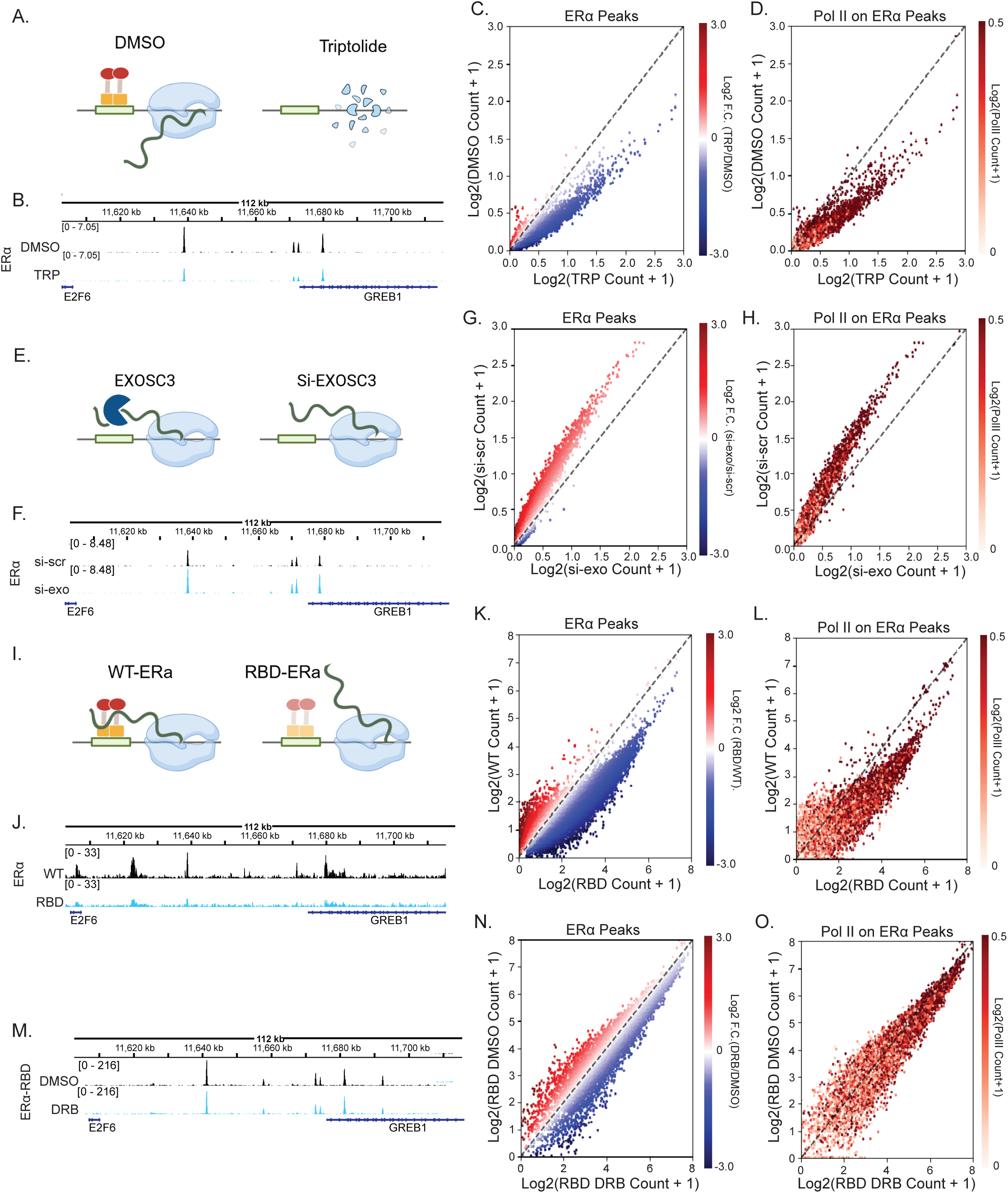
RNA from the paused sites contributes to TF accumulation. **A.** Schematic showing the mechanism of action of triptolide. **B.** IgV snapshot of ERα binding on GREB1 gene locus in DMSO and TRP. **C.** ERα signal on ERα peaks in DMSO and TRP. The y-axis denotes log2(CPM+1) values in DMSO, and the x-axis denotes log2(CPM+1) in TRP. The color of the spot denotes the log2F.C. (TRP/DMSO) value. **D.** Scatterplot plotted exactly like C. The color indicates the Pol II levels on those sites. **E.** Schematic showing the effect of exosc3 knockdown on ERα binding. **F.** IgV snapshot of ERα binding on GREB1 gene locus in scramble and exosc3 knockdown. **G.** ERα signal on ERα peaks in si-scr and si-exosc3. The y-axis denotes log2(CPM+1) values in si-scr, and the x-axis denotes log2(CPM+1) in si-exosc3. The color of the spot denotes the log2F.C. (si-exosc3/si-scr) value. **H.** Scatterplot plotted exactly like C. The color indicates the Pol II levels on those sites. **I.** Schematic showing the probable effect of RNA binding domain mutation on ERα binding. **J.** IgV snapshot of ERα binding on GREB1 gene locus in WT-ERα and RBD-ERα. **K.** ERα signal on ERα peaks in WT-ERα and RBD-ERα. The y-axis denotes log2(CPM+1) values in WT-ERα, and the x-axis denotes log2(CPM+1) in RBD-ERα. The color of the spot denotes the log2F.C. (RBD/WT) value. **L.** Scatterplot plotted exactly like C. The color indicates the Pol II levels on those sites. **M.** IgV snapshot of RBD-ERα binding on GREB1 gene locus in DMSO and DRB. **N.** ERα signal on RBD-ERα peaks in DMSO and DRB. The y-axis denotes log2(CPM+1) values in DMSO RBD-ERα, and the x-axis denotes log2(CPM+1) in DRB RBD-ERα. The color of the spot denotes the log2F.C. (DMSO/DRB) value. **O.** Scatterplot plotted exactly like C. The color indicates the Pol II levels on those sites.

Therefore, to test the involvement of chromatin-associated RNA in supplementing ERα binding, we increased its level RNA by knocking down the core subunit, exosc3 of the RNA exosome complex (Fig 4E). As opposed to DRB or TRP, RNA exosome degrades chromatin-associated RNA without affecting transcription *per se.* (Pefanis et al., 2015; Sigova et al., 2015). ChIP-seq of ERα upon exosc3 knockdown showed a gain in the binding for ERα, which was in line with the Pol II levels at the site (Fig 4F-H and S3B,E). The data suggests that chromatin-associated RNA can enhance the enrichment of ERα on chromatin.

ERα interacts with RNA through the canonical RNA binding domain (RRGG at aa 259-262 position) in its hinge region. The mutation of RRGG to AAAA abolishes RNA binding ability (Soota et al., 2024; Xu et al., 2021). We first tested the genomic binding of RNA binding deficient mutant (RBD) of ERα (RRGG to AAAA) without DRB. The RBD-ERα exhibited genome wide loss in binding with respect to WT (Fig 4I, S3E(Heatmaps)). Further, the highest loss was seen in the regions with the high level of Pol II (Fig 4J-L, and S3C, E).

With the presumption if RNA enriches ERα at paused sites by direct interaction, then the RBD-ERα should not gain binding upon DRB treatment. We performed ChIP-seq for the same and found indeed RBD-ERα did not show a gain of binding upon DRB treatment (Fig 4M-O and S3E). We further checked for the presence of nascent RNA at ERα gained sites upon DRB and found sites showing gain in ERα have higher nascent RNA upon DRB treatment (Fig S3G). Overall, the data suggests that the short RNA enriches ERα binding at Pol II paused sites through interaction with ERα.

### Polymerase-paused regions exhibit increased levels of H3K27ac

DRB treatment leads to increased promoter-proximal pausing, which is evident when we performed ChIP-seq for RNA Pol II after DRB treatment (Fig 5A). To check for DRB induced promoter pausing, we compared the gain with respect to the gene body and found it to be higher at the promoters (Fig S4A).

**Figure 5:**
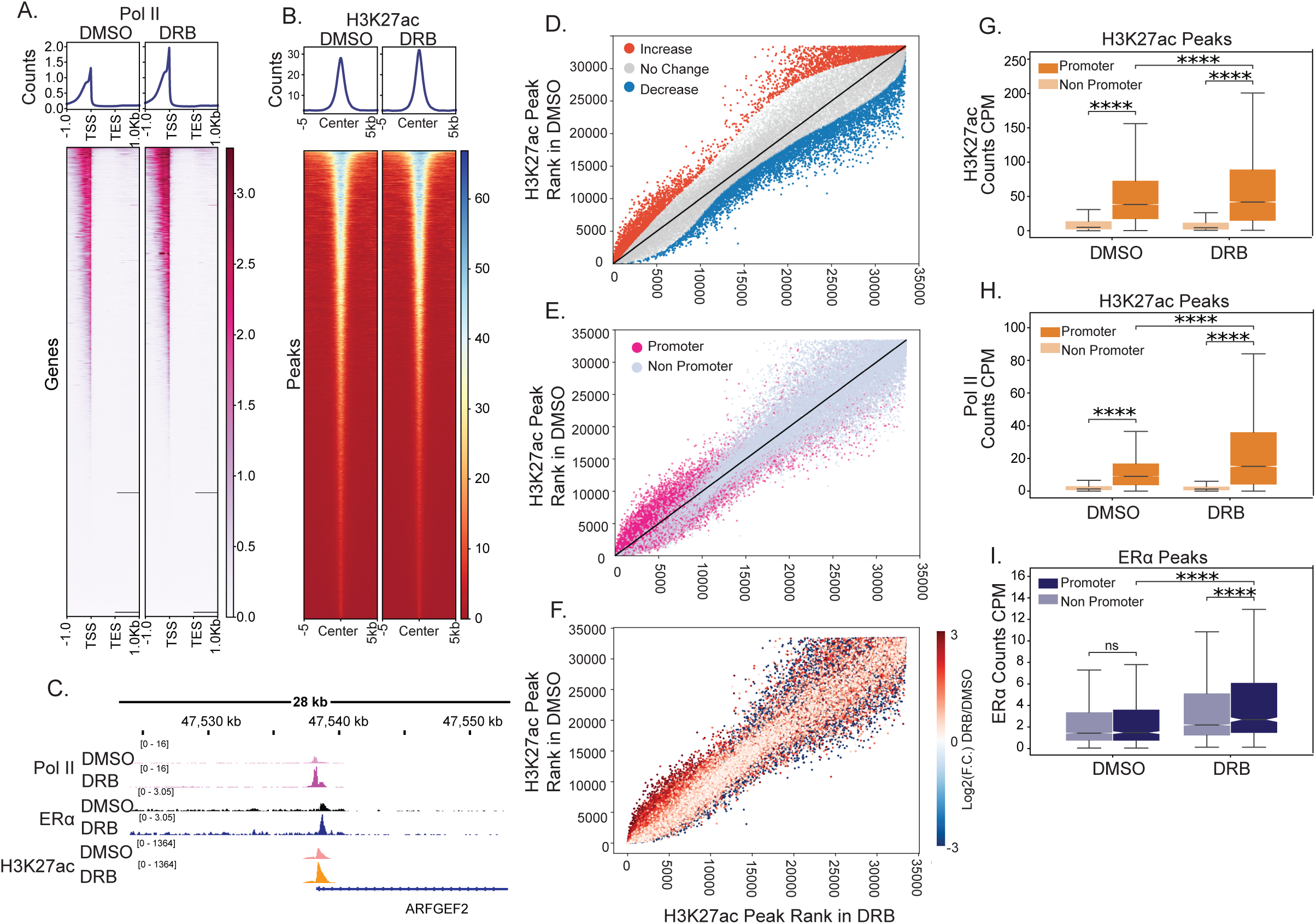
Polymerase-paused regions exhibit increased levels of H3K27ac. **A.** Heatmap of Pol II binding in DMSO and DRB at all expressing genes in MCF-7 cells. **B.** Heatmap of H3K27ac levels at H3K27ac peaks in DMSO and DRB. **C.** IgV snapshot of Pol II, ERα and H3K27ac binding on ARFGEF2 gene promoter in DMSO and DRB. **D.** H3K27ac peaks ranked on the basis of H3K27ac levels in DMSO on the y-axis and DRB on the x-axis. Red spots indicate regions showing an increase in H3K27ac upon DRB (Log2FC (DRB/DMSO > 0.5), Gray spots indicate regions showing no change in the levels (Fold change between −0.5 and 0.5), Blue spots indicate regions showing a decrease in the H3K27ac levels (Fold change less −0.5). **E.** Peak annotation of H3K27ac binding regions ranked as in **(G.)**, Pink denotes promoter regions, and Gray denotes non-promoter regions. **F.** Pol II gain on H3K27ac binding regions. The colour of the spot indicates the log2 fold change values post DRB treatment. **G.** Boxplot of H3K27ac levels on H3K27ac promoter and non-promoter regions in DMSO and DRB. p-values are calculated using the Mann-Whitney-Wilcoxon test for Promoter vs on-Promoter. p-values are calculated by the Wilcoxon signed-rank test for DMSO and DRB. **H.** Boxplot of Pol II levels on H3K27ac promoter and non-promoter regions in DMSO and DRB. p-values are calculated using the Mann-Whitney-Wilcoxon test for promoter vs non-promoter. p-values are calculated by the Wilcoxon signed-rank test for DMSO vs DRB. **I.** Boxplot of ERα levels on ERα promoter and non-promoter regions in DMSO and DRB. p-values are calculated using the Mann-Whitney-Wilcoxon test for promoter vs non-promoter. p-values are calculated by the Wilcoxon signed-rank test for DMSO and DRB.

We then asked if the gained ERα at paused Pol II regions, indeed is functional in creating transcriptional favourable environment. Towards this, we tested the levels of the H3K27ac as a proxy for transcriptional activity. A moderate gain in the levels of H3K27ac after DRB treatment was noted (Fig 5B-C). To understand what type of regions gained H3K27ac upon DRB, we first ranked the H3K27ac regions based on its levels in DRB and DMSO. Then, we categorized them as increasing, shown in Red (Log2FC(DRB/DMSO) > 0.5), non-changing, shown in Gray (−0.5 > Log2FC(DRB/DMSO) < 0.5) and decreasing, shown in Blue (Log2FC (DRB/DMSO) < −0.5) (Fig 5D). Gene annotation on the ranked regions was performed, which revealed that the regions showing a gain in H3K27ac were majorly promoters (Fig 5E and Fig S4B). Further, the RNA polymerase on the ranked regions was measured and found that H3K27ac peaks, which showed an increase, also exhibited increased polymerase pausing upon DRB treatment (Fig 5F).

We further quantified the levels of H3K27ac on promoters and non-promoter sites after DRB treatment and observed that these promoters already had higher levels of H3K27ac in DMSO. After DRB treatment, as polymerase gets paused at these sites, the H3K27ac levels increased further (Fig 5G-H, S4C-F). Intriguingly, the levels of ERα were similar in promoter and non-promoter regions. However, the ERα binding on the promoter showed a higher gain due to polymerase pausing after DRB (Fig 5I). In addition to paused polymerase, ERα promoters showed increased H3K27ac levels as well (Fig S4G). Overall, the data suggests that the paused polymerase stabilizes the binding of ERα on the promoters, and these promoters also show a gain of H3K27ac levels suggestive of an enhanced transcriptional environment at these regions as compared to the non-paused state.

### Higher ERα binding and H3K27ac gain upon Pol II pausing correlates with increased transcription after pausing release

In order to test if the favourable environment created by ERα at paused sites indeed allows more transcription of genes once pausing is released. We analysed nascent transcripts from BrU-seq after 10 mins of release from DRB treatment and tested the transcription levels in region 2kb downstream of the promoter (Veloso et al., 2014). We binned the promoter regions based on increased transcription after release vs. without DRB treatment (Fig 6A-B). The promoters that showed the highest transcription after the release from DRB were the ones that also exhibited a higher gain of H3K27ac and ERα upon DRB treatment (Fig 6C-E, S5A-B). The data suggests that the gain in ERα binding and H3K27ac at paused promoters facilitated more robust transcription after pausing was released (Fig 6F).

**Figure 6:**
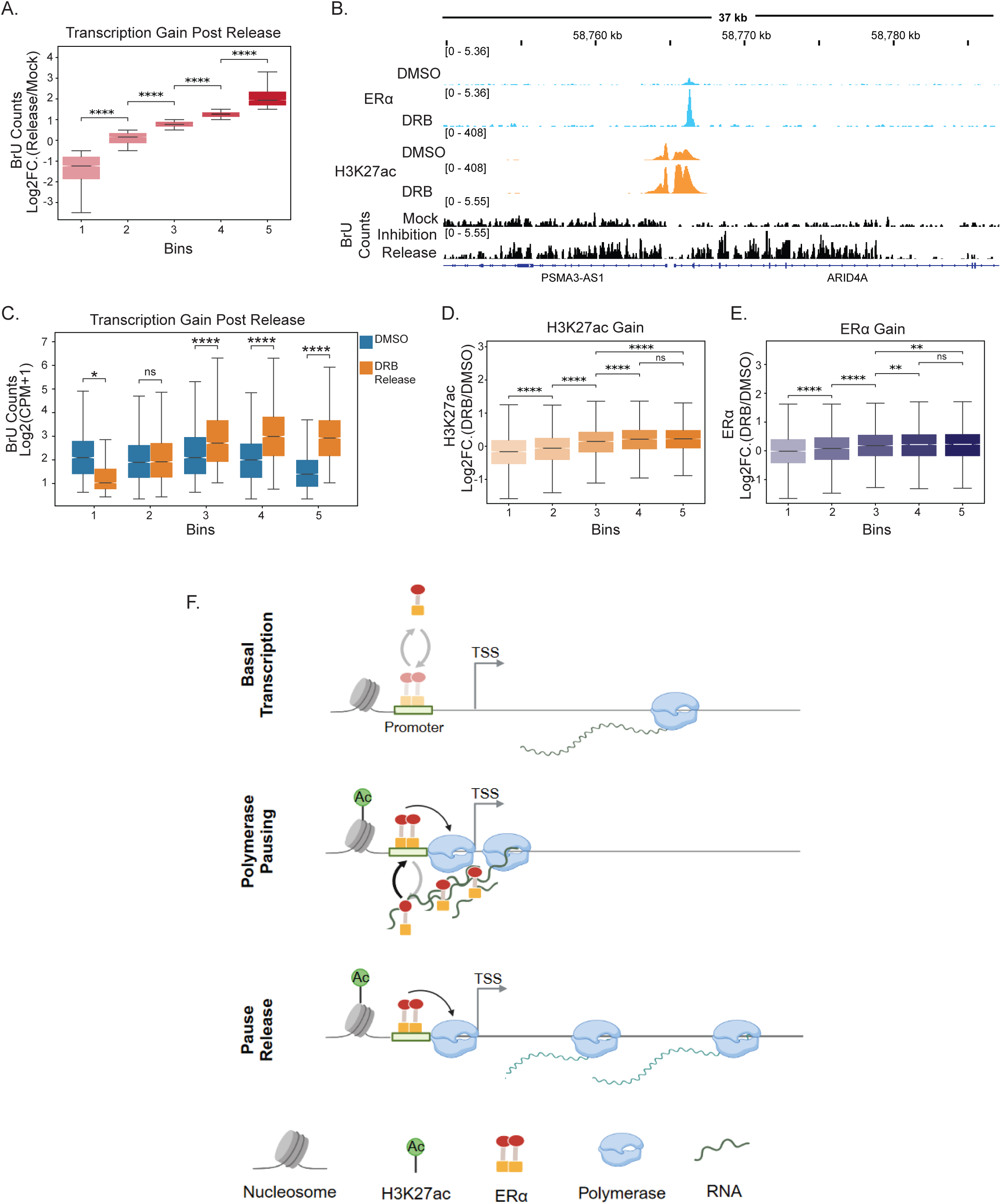
Higher ERα binding and H3K27ac gain upon Pol II pausing correlates with increased transcription after pausing release. A. Log2F.C. (DRB Release/DMSO) of normalized BrU counts. In the categories based on promoter firing, i.e. BrU counts from (2kb downstream from TSS) upon release. 1 = Fold Change < −0.5, 2 = −0.5 < Fold Change < 0.5, 3 = 0.5 < Fold Change < 1.0, 4 = 1.0 < Fold Change < 1.5, 5 = Fold Change > 0.5. **B.** IgV snapshot of ERα and H3K27ac levels in DMSO and DRB on ARID4A locus and BrU levels in DMSO and after release **C.** BrU levels in DMSO and DRB in the categories. **D. and E.** Log2FC(DRB/DMSO) for H3K27ac and ERα levels, respectively, on promoter categories from **A.**

## Discussion

Promoter-proximal pausing is a key regulatory step in transcription. Typically, RNA polymerase pauses within 20-60 nucleotides of the transcription start site (Core et al., 2008; Muse et al., 2007; Zeitlinger et al., 2007). We treated cells with DRB, an elongation inhibitor that induces polymerase pausing, and observed a significant increase in ERα binding across the genome (Fig 1B-C). In contrast, TRP, an initiation inhibitor, caused a loss of ERα binding. The increased binding with DRB treatment occurred primarily at sites with higher Pol II levels, predominantly at promoters (Fig 1H). Using SMT tracking, we found that both the binding fraction and dwell time of ERα increased upon DRB treatment (Fig 3), which we attribute to ERα’s RNA-binding ability. This stable ERα binding correlated with increased H3K27ac levels, suggesting enhanced transcription of paused genes upon release from pausing.

The BAF complex dynamically creates unstable nucleosomes. Paused polymerase recruits BAF to these paused sites, where the presence of transcription factors (TFs) may stabilize the BAF, leading to nucleosome eviction and the formation of nucleosome-free regions (NFRs) (Brahma & Henikoff, 2024). Our data showed that paused sites recruit ERα upon estrogen stimulation, and this binding was further enhanced by DRB exposure. We propose that the stable binding and prolonged dwell time of ERα during elongation inhibition are critical for recruiting BAF complexes to stabilize NFRs.

Although elongation inhibitors reduce overall RNA levels, short RNAs are still transcribed from paused polymerase (Henriques et al., 2013; Veloso et al., 2014). We found that the sites that showed ERα and H3K27ac gain also had higher RNA levels in the presence of DRB (Fig S3F, G). This stable chromatin-bound ERα fraction likely results from physical interactions between ERα and short RNAs produced at these paused sites. Supporting this, an ERα mutant that cannot bind RNA did not show gain as observed with WT. Thus, RNA binding may reduce the strict requirement for strong TF motifs at these sites (Soota et al., 2024). In summary, transcription factor at paused sites is stabilized through their RNA-binding ability.

Our data suggest that TF binding near promoter-proximal pausing creates a favourable environment for robust transcription by priming these sites with active machinery. H3K27ac, a marker of active chromatin, increased after DRB treatment, particularly at promoter sites with polymerase pausing (Fig 5E-F). Upon release from DRB inhibition, these promoters showed significantly higher transcription, indicating heightened transcriptional potential post-pausing (Fig 6). These findings suggest that promoters may use Pol II pausing to enhance their gene transcription by directly recruiting transcription factor complexes in RNA-dependent manner.

## Materials and Methods

### Cell Culture

MCF-7 cells were obtained from ATCC and were cultured in High glucose DMEM media (11965092, Invitrogen) with 10% FBS (10437028, Invitrogen). Cells were grown at 37°C and 5% CO_2_ conditions in an incubator.

### Ligand stimulation and transcription inhibitor treatment

For experiments involving E2 stimulations, cells were seeded on Day 0. After 24 hours, cells were washed with DPBS (Invitrogen, 14190144), and the media was changed to stripping media (high glucose DMEM without phenol red (<u>1063029</u>, Invitrogen) and 5% charcoal-stripped FBS (12676029, Invitrogen)). Cells were kept in stripping media for three days. After 72 hours, cells were treated with β-estradiol (E2758, Sigma-Aldrich) at a concentration of 10nM for 1 hour.

In the case of experiments involving transcription inhibitor treatment, cells were treated with DRB (D1916, Sigma) at 10uM and TRP (T3652, Sigma) at 1uM or DMSO for 1 hour. After 1 hour of inhibitor treatment, 10nM of E2 was added in the media and processed for Immunostaining and ChIP-seq. In case of live cell imaging, cells were imaged as soon as E2 was added, till 1hour of treatment was completed.

### Transfections

For plasmid transfection, the cells were seeded and kept in stripping media for 48 hours. After 48 hours of stripping 1μg of ERα-eGFP/Halo plasmid was transfected using OptiMEM without phenol red with lipofectamine 3000. Cells are processed after 24 hours of transfections.

For siRNA transfections, two rounds of transfection were done. First transfection was done 24hours after cell seeding. Second round of transfection was done after 48hours of keeping cells in stripping media.

### ChIP-seq

MCF-7 cells were grown as mentioned above. They were hormone-stripped for 72 hours. The cells were treated with inhibitors as mentioned above. After two hours of inhibitor treatment and 1 hour of E2 treatment. Cells were fixed using 1% formaldehyde for 10 mins at room temperature on an orbital shaker. After 10mins, formaldehyde was quenched using 0.125M glycine for 5mins at room temperature on the orbital shaker. The cells were washed with cold 1XPBS and scraped. The cells were pelleted down at 3000rpm for 5mins at 4°C. The pellets were stored at −80°C. 10 million cells were resuspended in 1 ml of nuclear lysis buffer (NLB) (50mM Tris-HCl pH 7.4, 1% SDS, 10mM EDTA pH 8.0, and 1X PIC) and incubated in ice for 10mins. After nuclear lysis, cells were sonicated using Bioruptor pico for 28cycles with 30secs ON and 30secs OFF settings. The lysate was then spun at 12000rpm for 12mins at 4°C to clear off the cell debris. 100μg of chromatin was used for IP, further 3μg of sonicated S2 chromatin was added as spike-in for only H3K27ac ChIP. The IP was diluted 2.5 times with dilution buffer (20mM Tris-HCl pH 7.4, 100mM NaCl, 2mM EDTA pH 8.0, 0.5% Triton X-10, and 1X PIC). IP was set in a volume of 500μl. Additionally, 50μl was taken out and labelled as 10% input. For IP, main as well spike-in antibody was added in their respective amounts (ERα (sc-8002, Santa Cruz Biotech) 1μg, anti-Pol II (Diagenode C15200004) 2μg, H3K27ac (ab4729, Abcam) 1μg and spike-in H2av 0.3ug (61686, Active Motif)). IP was incubated with antibodies overnight at 4°C at constant rotation of 12rpm. After incubation, 15μl of Protein G beads pre-blocked in BSA were added to the mix. It was incubated for 4 hours at 4°C on the rotor. After incubation, the beads were washed with wash buffers I, II, III and 1XTE sequentially (Li et al., 2013). The DNA-protein complexes were eluted from the beads by resuspending the beads in (100mM NaHCO3, 1%SDS, 2μg of RNaseA) at 37°C in a thermomixer with 1400rpm for 30mins. The crosslinked DNA-protein complexes were reverse-crosslinked over night at 65°C by the addition of NaCl. After that 2μl of Proteinase K (20mg/ml) was added to the mix and incubated for 2hours at 45°C. The ChIP DNA was then purified using PCI (Phenol:Chloroform:Isoamylalcohol) and ethanol precipitation. The pellet was resuspended in 10μl of NFW and used for library preparation using NEBNext Ultra II DNA library Prep kit with Sample Purification Beads (E7103L). The sequencing was performed on HiSeq/Nova-seq platform.

### DRB-BrU Seq Analysis

Raw files were first aligned with hg19 genome assembly using bowtie2 with default options (Langmead & Salzberg, 2012; Zhang et al., 2008). rRNA were separately aligned for normalization control. Reads with quality score less than 30 were filtered out. The reads were first down sampled according to the sample with the lowest sequencing depth. Then, reads were further down sampled according to rRNA reads.

### Alignment and Spike-in normalization

Raw fastq reads were aligned to the hg19 genome for human and dm3 reference genome for drosophila. Reads with quality score less than 30 were filtered out. The reads were first down sampled according to the sample with the lowest sequencing depth. Then, reads were further down sampled according to spike-in factor calculated as shown in (Egan et al., 2016).

### Generation of Plasmids

EFS-ERα-Halo was constructed in PX459 plasmid using golden gate assembly. EFS, ERα and Halo were separately amplified from pHAGE-EFS-N22p-3XRFPnls (Addgene: 75387), pEGFP-C1-ER alpha (Addgene: 28230) and pcDNA5-H2B_Halo_T2A_EGFP (Addgene:135444) respectively (Frei et al., 2019; Ma et al., 2016; Stenoien et al., 2000). All the fragments were then subjected to golden gate assembly with BsaI-v2 (R3733L, NEB).

### Live Cell Imaging

Cells were transfected with ERα-eGFP following the protocol mentioned above. For live cell imaging cells were imaged using PLAPON 60x/1.42 oil objective of Olympus FV3000 microscope. Microscope is equipped with incubator to keep temperature at 37°C and 5% CO_2_. The frame rate was kept at 15 sec/frame (10.6s camera time and 4.4s rest time). The cells were imaged for 1hour for a total of 240 frames.

### Immunostaining

For experiments involving removal of nucleoplasmic proteins. Cells were first treated with CSK buffer for 3mins at RT (PIPES/KOH 10mM pH6.8, NaCl 100mM, Sucrose 300mM, EGTA 1mM, MgCl2 1mM, DTT 1mM, 1XPIC, 0.5% TritonX100. After treatment cells are immediately fixed with 4%PFA in 1XPBS for 15mins. In other experiments cells were directly fixed with 4%PFA in 1XPBS for 15mins. Fixing is followed by 3 washes with 1XPBS. Cells are then permeabilised and blocked with 1% BSA with 0.2% TritonX100 in PBS). After which cells are incubated with primary antibody (1μg/ml ERα, YY1 and Pol IIS5P) at 4°C overnight. It was followed by 3 washes with 1XPBST. Then, cells are incubated with appropriate secondary antibody in 1/500 dilution for 1hour at RT. It was followed by 3 washes with 1XPBST. Cells were then incubated with Hoechst 3342 for 10mins at RT and then mounted in 90% glycerol.

### Fixed cell imaging

Cells were imaged in LSM980 with confocal module using 63X oil objective. In experiments (Supp 2A and B) images were acquired using airyscan2 detector.

### Single-molecule imaging and tracking

The MCF-7 cells were transiently transfected with ERα-HaloTag plasmid. This plasmid expresses ERalpha-HaloTag from the EFS promoter. After transfection, the cells were treated with 5 nM JF646 HaloTag Ligand (HTL) (Grimm et al., 2015) for 20 min. The cells were then washed thrice for 15 min with stripping media to remove the unbound HaloTag ligand. Subsequently, the cells were treated with either 10 µM DRB or DMSO for 120 mins. The cells were then imaged for ∼1 hr at 37 °C in 5% CO_2_ using a Leica DMi8 infinity TIRF inverted fluorescence microscope equipped with a Photometric Prime95B CMOS camera, 100X 1.47 NA TIRF objective lens, 638nm 150 mW laser module. Time-lapse movies were acquired with two imaging regimes: 1) fast imaging regime (15 ms time interval (10 ms exposure + 5 ms camera processing time), 100% laser power, 1000 frames) and 2) slow imaging regime (200 ms time interval (50 ms exposure + 150 ms interval), 30% laser power, 600 frames).

For quantifying diffusion parameters (fraction of bound/unbound molecules), the single-molecule tracking was performed using movies acquired with fast imaging regime using DiaTrack Version 3.05 (Vallotton and Olivier 2013), with the following settings as previously described (Podh et al., 2022; Ranjan et al., 2020): remove blur: 0.07; remove dim: 45-150; maximum jump: 6 pixels where each pixel was 110 nm. This software determines the precise position of single molecules by Gaussian intensity fitting and assembles particle trajectories over multiple frames. The trajectory data exported from Diatrack was further converged into a single .csv file using a custom computational package ‘Sojourner’ (https://rdrr.io/github/sheng-liu/sojourner/). The Spot-On analysis was performed on three frames or longer trajectories using the web interface https://spoton.berkeley.edu/ (Hansen et al., 2018). The bound fractions and diffusion coefficients were extracted from the CDF of observed displacements over different time intervals. The cumulative displacement histograms were fitted with a 2-state model.

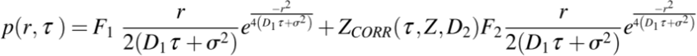

Where F_1_ and F_2_ are bound and free fractions, σ is single molecule localization error, D_1_ and D_2_ are diffusion coefficients of bound and free fractions, and Z_CORR_ is the correction factor for fast molecules moving out of axial detection range (Hansen et al., 2018). The following settings were used on the Spot-On web interface: bin width: 0.01 µm; number of time points: 8: jumps to consider: 4 pixels; use entire trajectories: No: max jump: 1 µm. For model fitting the following parameters were selected: D_bound_ (µm^2^/s): min 0.0001 max 0.5, D_free_ (µm^2^/s): min 0.5 max 5, F_bound_: min 0 max 1, Localization error (µm): Fit from data: Yes (min 0.01 max 0.1); dZ (µm): 0.65 for JF646: Use Z Correction: Yes; Model Fit: CDF; Iterations: 3.

For residence time analysis, the single-molecule tracking was performed using movies acquired with slow imaging regime using the “TrackRecord” software developed in Matlab (The Matworks Inc.) (Mazza et al., 2013). The software provides automated features for particle detection (using intensity thresholding), tracking (using the nearest neighbour algorithm with molecules allowed to move a maximum of 6 pixels from 1 frame to the next, and only tracks that are at least 4 frames or longer are kept. Gaps to close 4 frames to compensate for fluorophore blinking), photobleaching correction, and quantification of residence time. The residence time was determined by the fitting to the survival distribution. Briefly, the survival histogram was generated from the time periods that each molecule is stationary. In practice, even tightly bound particles move slightly due to chromatin and nuclear motion, and therefore a maximum frame-to-frame displacement of 262 nm (R_min_), and a two-frame displacement of 370 nm (R_max_) (both obtained from the motion of the chromatin-bound histone H2B) have been used to define bound portions of each particle’s track. Because there is a chance that even a fast-diffusing molecule will move less than these thresholds, a further constraint on the minimum number of time points in the bound segment for each particle (N_min_) is used to reduce <1% the contribution of diffusing molecules to the survival histogram. The N_min_ value used for 200 ms time-interval movies was 4. The total bound fraction is then calculated as the ratio of bound track segments to the total number of particles. To extract residence times, the survival distribution, S(t), is fit by least squares to a mixed exponential decay with two rate constants, *k*_ns_=1/T_ns_ and *k*_s_ = 1/T_s_:

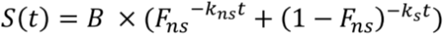

Where B is the bound fraction, and Fns is the fraction of particles non-specifically bound. To check for over-fitting, the distribution is also fit to a single-component exponential:

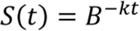

The fits are compared using an F-test to ensure that the two-component model gives a significant improvement over the single-component decay. Z-test was performed

### Image Analysis

Image analysis was done using Matlab. ERα spots were called differently based on the type of imaging-

1. Live Cell Images: The nucleus was segmented from the ERα signal. The nucleus was first blurred using gaussian blur. Then, the blurred image was subjected to Otsu thresholding. For spot calling, we utilized unsharp masking. blurred image was subtracted from the original image to get high intensity regions. Then masks of high difference regions were created. The spots were then subjected to size thresholding where regions below 6 pixels were removed. For calculating number density: Number of the spots at every frame were divided by the area of the nucleus at that frame. For calculating average diameter: Diameter of all the spots were calculated by taking the max axis length. Then average of all the diameter at a given frame was calculated. The raw curves from both the average diameter and number density with time were subjected to moving average filter with the kernel size of 10. The maxima was then calculated from the filtered curve. The t-half was calculated after normalizing the curve in min-max fashion followed by fitting exponential equation to get the rate. Then the rate was used to calculate the t-half.
2. Fixed Cell Images: Otsu thresholding was used to segment the nucleus from the Hoechst channel. For calling spots first high boost filtering was used to segment high intensity spots. Then size thresholding was done to random blips. Further watershed algorithm was used to segment close spots. Volume fraction was calculated getting the sum of volume of all the spots divided by the volume of the nucleus.

## Contributions

RM and DN conceived the project. RM performed and analysed most experiments with help from DS. AD, NKP and GM performed the SMT experiments and analysed SMT data. RM and DN wrote the manuscripts with input from all the authors.

## Acknowledgement

This work was supported by the Department of Atomic Energy, Government of India, Project Identification No. RTI 4006. DN acknowledges funding support from India Alliance-Wellcome trust (IA/1/14/2/501539). RM and DS were supported by TIFR-NCBS graduate program. GM acknowledges funding support from the Department of Biotechnology, Govt. of India (BT/INF/22/SP53103/2024). We thank Awadhesh Pandit for technical help in NGS. We acknowledge technical support from Genomics and Central Imaging facility at NCBS.

## Legends

**Figure S1 (Related to Figure 1) A.**

Expression levels of ESR1, Myc and Pre-Myc gene after treating the cells with DMSO (mock), DRB and TRP. p-values are calculated using an unpaired t-test. **B.** Boxplot of motif score on the ERα peaks sorted based on different categories. p-values are calculated using the Mann-Whitney-Wilcoxon test. **C.** Boxplot showing ERα levels in DMSO and DRB on ERα peaks in the categories sorted based on motif and Pol II levels. p-values are calculated using the Mann-Whitney-Wilcoxon test.

**Figure S2 (Related to Figure 2) A.**

Confocal images of MCF-7 cells immunostained with ERα and Pol IIS5P at 60, 120 and 180 mins post E2 addition. Cells were treated with ICI for 24 hours. **B. and C.** Volume Fraction and Normalized Overlap for ERα condensate and Pol IIS5P spots at different time points of E2 signalling. p-values are calculated using the Mann-Whitney-Wilcoxon test. **D.** Confocal images of MCF-7 cells immunostained with Erα and Nascent RNA (BrU) at 60, 120 and 180 mins post E2 addition. **E.** Normalized Overlap for ERα condensate and BrU spots at different time points of E2 signalling. p-values are calculated using the Mann-Whitney-Wilcoxon test. **F.** Scatter plot of nuclear mean intensity on the y-axis and max number density on the x-axis. Pearson correlation coefficient of 0.77. The black line denotes the line of best fit. **G.** Scatter plot of nuclear mean intensity on the y-axis and t-half calculated from number density curve on the x-axis. Pearson correlation coefficient of −0.48. The black line denotes the line of best fit. **H.** Scatter plot of nuclear mean intensity on the y-axis and max average diameter on the x-axis. Pearson correlation coefficient of 0.42. The black line denotes the line of best fit. **I.** Scatter plot of nuclear mean intensity on the y-axis and t-half calculated from average diameter curve on the x-axis. Pearson correlation coefficient of 0.01. The black line denotes the line of best fit. **J.** Boxplot of mean intensity of ERα-eGFP at the first frame in DMSO and DRB conditions. p-values are calculated using the Mann-Whitney-Wilcoxon test.

**Figure S3 (Related to Figure 4) A.**

ERα binding loss quantified upon triptolide treatment Log2FC(TRP/DMSO) in increasing Pol II level bins. **B.** ERα binding gain quantified upon exosc3 knockdown Log2FC (si-exo/si-scr) in increasing Pol II level bins. **C.** ERα binding loss quantified in RBD-ERα w.r.t WT-ERα Log2FC(RRGG/WT) in increasing Pol II level bins. **D.** ERα binding quantified in DMSO RBD-ERα w.r.t DRB RBD-ERα Log2FC(DRB/DMSO) in increasing Pol II level bins. **E.** Heatmap for the peaks in Fig 4C, G, K, O. **F. and G.** 4SU signal log2(CPM+1) in DRB on regions showing increase, decrease and no-change in binding of H3K27ac, ERα upon treating cells with DRB.

**Figure S4 (Related to Figure 5) A.**

Boxplot of Pol II gain on promoter and gene body upon DRB treatment. **B.** H3K27ac peaks ranked on the basis of H3K27ac levels in DMSO on the y-axis and DRB on the x-axis. Color denotes the annotation of H3K27ac. Pink, light green, orange, purple, dark green and yellow denote promoter, distal intergenic, 3’ UTR, exon, intron, and 5’ UTR, respectively. **C.** Boxplot of top 1500 non-changing (log2 fold change −0.5 to 0.5) and increasing (log2 fold change 0.5) H3K27ac regions. The y-axis represents the log2F.C.(DRB/DMSO) for H3K27ac and H3K27ac levels (CPM) in DMSO and DRB. **D.** Boxplot of top 1500 non-changing (log2 fold change −0.5 to 0.5) and increasing (log2 fold change 0.5) H3K27ac regions. The y-axis represents the log2F.C.(DRB/DMSO) for Pol II and Pol II levels (CPM) in DMSO and DRB. **E.** H3K27ac levels in increasing Pol II level bins. **F.** H3K27ac binding gain quantified upon DRB treatment Log2F.C.(DRB/DMSO) in increasing Pol II level bins. **G.** Heatmap showing H3K27ac and Pol II levels on promoter and non-promoter ERα peaks in DMSO and DRB.

**Figure S5 (Related to Figure 6)**

**A. and B.** H3K27ac and ERα levels in DMSO and DRB from promoter categories in 6A.

## References

1. Adelman, K., & Lis, J. T. (2012). Promoter-proximal pausing of RNA polymerase II: Emerging roles in metazoans. Nature Reviews. Genetics, 13(10), 720–731. 10.1038/nrg3293

2. Aibara, S., Schilbach, S., & Cramer, P. (2021). Structures of mammalian RNA polymerase II pre-initiation complexes. Nature, 594(7861), 124–128. 10.1038/s41586-021-03554-8

3. Bartman, C. R., Hamagami, N., Keller, C. A., Giardine, B., Hardison, R. C., Blobel, G. A., & Raj, A. (2019). Transcriptional Burst Initiation and Polymerase Pause Release Are Key Control Points of Transcriptional Regulation. Molecular Cell, 73(3), 519–532.e4. 10.1016/j.molcel.2018.11.004

4. Beckman, W. F., Jiménez, M. Á. L., Moerland, P. D., Westerhoff, H. V., & Verschure, P. J. (2021). 4sUDRB-sequencing for genome-wide transcription bursting quantification in breast cancer cells (p. 2020.12.23.424175). bioRxiv. 10.1101/2020.12.23.424175

5. Brahma, S., & Henikoff, S. (2024). The BAF chromatin remodeler synergizes with RNA polymerase II and transcription factors to evict nucleosomes. Nature Genetics, 56(1), 100–111. 10.1038/s41588-023-01603-8

6. Core, L., & Adelman, K. (2019). Promoter-proximal pausing of RNA polymerase II: A nexus of gene regulation. Genes & Development, 33(15–16), 960–982. 10.1101/gad.325142.119

7. Core, L. J., Waterfall, J. J., & Lis, J. T. (2008). Nascent RNA Sequencing Reveals Widespread Pausing and Divergent Initiation at Human Promoters. Science, 322(5909), 1845–1848. 10.1126/science.1162228

8. Dror, I., Golan, T., Levy, C., Rohs, R., & Mandel-Gutfreund, Y. (2015). A widespread role of the motif environment in transcription factor binding across diverse protein families. Genome Research, 25(9), 1268–1280. 10.1101/gr.184671.114

9. Egan, B., Yuan, C.-C., Craske, M. L., Labhart, P., Guler, G. D., Arnott, D., Maile, T. M., Busby, J., Henry, C., Kelly, T. K., Tindell, C. A., Jhunjhunwala, S., Zhao, F., Hatton, C., Bryant, B. M., Classon, M., & Trojer, P. (2016). An Alternative Approach to ChIP-Seq Normalization Enables Detection of Genome-Wide Changes in Histone H3 Lysine 27 Trimethylation upon EZH2 Inhibition. PLOS ONE, 11(11), e0166438. 10.1371/journal.pone.0166438

10. Falo-Sanjuan, J., Lammers, N. C., Garcia, H. G., & Bray, S. J. (2019). Enhancer Priming Enables Fast and Sustained Transcriptional Responses to Notch Signaling. Developmental Cell, 50(4), 411–425.e8. 10.1016/j.devcel.2019.07.002

11. Frei, M. S., Hoess, P., Lampe, M., Nijmeijer, B., Kueblbeck, M., Ellenberg, J., Wadepohl, H., Ries, J., Pitsch, S., Reymond, L., & Johnsson, K. (2019). Photoactivation of silicon rhodamines via a light-induced protonation. Nature Communications, 10(1), 4580. 10.1038/s41467-019-12480-3

12. Gilchrist, D. A., Nechaev, S., Lee, C., Ghosh, S. K. B., Collins, J. B., Li, L., Gilmour, D. S., & Adelman, K. (2008). NELF-mediated pausing of Pol II can enhance gene expression by blocking promoter-proximal nucleosome assembly. Genes & Development, 22(14), 1921–1933. 10.1101/gad.1643208

13. Gilchrist, D. A., Santos, G. D., Fargo, D. C., Xie, B., Gao, Y., Li, L., & Adelman, K. (2010). Pausing of RNA Polymerase II Disrupts DNA-Specified Nucleosome Organization to Enable Precise Gene Regulation. Cell, 143(4), 540–551. 10.1016/j.cell.2010.10.004

14. Gressel, S., Schwalb, B., & Cramer, P. (2019). The pause-initiation limit restricts transcription activation in human cells. Nature Communications, 10(1), 3603. 10.1038/s41467-019-11536-8

15. Grimm, J. B., English, B. P., Chen, J., Slaughter, J. P., Zhang, Z., Revyakin, A., Patel, R., Macklin, J. J., Normanno, D., Singer, R. H., Lionnet, T., & Lavis, L. D. (2015). A general method to improve fluorophores for live-cell and single-molecule microscopy. Nature Methods, 12(3), 244–250. 10.1038/nmeth.3256

16. Gu, B., Comerci, C. J., McCarthy, D. G., Saurabh, S., Moerner, W. E., & Wysocka, J. (2020). Opposing Effects of Cohesin and Transcription on CTCF Organization Revealed by Super-resolution Imaging. Molecular Cell, 80(4), 699–711.e7. 10.1016/j.molcel.2020.10.001

17. Hah, N., Danko, C. G., Core, L., Waterfall, J. J., Siepel, A., Lis, J. T., & Kraus, W. L. (2011). A rapid, extensive, and transient transcriptional response to estrogen signaling in breast cancer cells. Cell, 145(4), 622–634. 10.1016/j.cell.2011.03.042

18. Hansen, A. S., Woringer, M., Grimm, J. B., Lavis, L. D., Tjian, R., & Darzacq, X. (2018). Robust model-based analysis of single-particle tracking experiments with Spot-On. eLife, 7, e33125. 10.7554/eLife.33125

19. Henriques, T., Gilchrist, D. A., Nechaev, S., Bern, M., Muse, G. W., Burkholder, A., Fargo, D. C., & Adelman, K. (2013). Stable Pausing by RNA Polymerase II Provides an Opportunity to Target and Integrate Regulatory Signals. Molecular Cell, 52(4), 517–528. 10.1016/j.molcel.2013.10.001

20. Inukai, S., Kock, K. H., & Bulyk, M. L. (2017). Transcription factor–DNA binding: Beyond binding site motifs. Current Opinion in Genetics & Development, 43, 110–119. 10.1016/j.gde.2017.02.007

21. Keenan, S. E., Blythe, S. A., Marmion, R. A., Djabrayan, N. J.-V., Wieschaus, E. F., & Shvartsman, S. Y. (2020). Rapid dynamics of signal-dependent transcriptional repression by Capicua. Developmental Cell, 52(6), 794. 10.1016/j.devcel.2020.02.004

22. Langmead, B., & Salzberg, S. L. (2012). Fast gapped-read alignment with Bowtie 2. Nature Methods, 9(4), 357–359. 10.1038/nmeth.1923

23. Li, W., Notani, D., Ma, Q., Tanasa, B., Nunez, E., Chen, A. Y., Merkurjev, D., Zhang, J., Ohgi, K., Song, X., Oh, S., Kim, H.-S., Glass, C. K., & Rosenfeld, M. G. (2013). Functional roles of enhancer RNAs for oestrogen-dependent transcriptional activation. Nature, 498(7455), 516–520. 10.1038/nature12210

24. Liu, L., Xu, Y., He, M., Zhang, M., Cui, F., Lu, L., Yao, M., Tian, W., Benda, C., Zhuang, Q., Huang, Z., Li, W., Li, X., Zhao, P., Fan, W., Luo, Z., Li, Y., Wu, Y., Hutchins, A. P., … Esteban, M. A. (2014). Transcriptional pause release is a rate-limiting step for somatic cell reprogramming. Cell Stem Cell, 15(5), 574–588. 10.1016/j.stem.2014.09.018

25. Ma, H., Tu, L.-C., Naseri, A., Huisman, M., Zhang, S., Grunwald, D., & Pederson, T. (2016). Multiplexed labeling of genomic loci with dCas9 and engineered sgRNAs using CRISPRainbow. Nature Biotechnology, 34(5), 528–530. 10.1038/nbt.3526

26. Mann, R., & Notani, D. (2023). Transcription factor condensates and signaling driven transcription. Nucleus, 14(1), 2205758. 10.1080/19491034.2023.2205758

27. Materne, P., Anandhakumar, J., Migeot, V., Soriano, I., Yague-Sanz, C., Hidalgo, E., Mignion, C., Quintales, L., Antequera, F., & Hermand, D. (2015). Promoter nucleosome dynamics regulated by signalling through the CTD code. eLife, 4, e09008. 10.7554/eLife.09008

28. Mazza, D., Abernathy, A., Golob, N., Morisaki, T., & McNally, J. G. (2012). A benchmark for chromatin binding measurements in live cells. Nucleic Acids Research, 40(15), e119. 10.1093/nar/gks701

29. Muse, G. W., Gilchrist, D. A., Nechaev, S., Shah, R., Parker, J. S., Grissom, S. F., Zeitlinger, J., & Adelman, K. (2007). RNA polymerase is poised for activation across the genome. Nature Genetics, 39(12), 1507–1511. 10.1038/ng.2007.21

30. Nair, S. J., Yang, L., Meluzzi, D., Oh, S., Yang, F., Friedman, M. J., Wang, S., Suter, T., Alshareedah, I., Gamliel, A., Ma, Q., Zhang, J., Hu, Y., Tan, Y., Ohgi, K. A., Jayani, R. S., Banerjee, P. R., Aggarwal, A. K., & Rosenfeld, M. G. (2019). Phase separation of ligand-activated enhancers licenses cooperative chromosomal enhancer assembly. Nature Structural & Molecular Biology, 26(3), 193–203. 10.1038/s41594-019-0190-5

31. Nechaev, S., Fargo, D. C., dos Santos, G., Liu, L., Gao, Y., & Adelman, K. (2010). Global Analysis of Short RNAs Reveals Widespread Promoter-Proximal Pausing and Arrest of Pol II in Drosophila. Science, 327(5963), 335–338. 10.1126/science.1181421

32. Oksuz, O., Henninger, J. E., Warneford-Thomson, R., Zheng, M. M., Erb, H., Vancura, A., Overholt, K. J., Hawken, S. W., Banani, S. F., Lauman, R., Reich, L. N., Robertson, A. L., Hannett, N. M., Lee, T. I., Zon, L. I., Bonasio, R., & Young, R. A. (2023). Transcription factors interact with RNA to regulate genes. Molecular Cell, 83(14), 2449–2463.e13. 10.1016/j.molcel.2023.06.012

33. Pefanis, E., Wang, J., Rothschild, G., Lim, J., Kazadi, D., Sun, J., Federation, A., Chao, J., Elliott, O., Liu, Z.-P., Economides, A. N., Bradner, J. E., Rabadan, R., & Basu, U. (2015). RNA Exosome-Regulated Long Non-Coding RNA Transcription Controls Super-Enhancer Activity. Cell, 161(4), 774–789. 10.1016/j.cell.2015.04.034

34. Podh, N. K., Das, A., Dey, P., Paliwal, S., & Mehta, G. (2022). Single-molecule tracking for studying protein dynamics and target-search mechanism in live cells of *S. cerevisiae*. STAR Protocols, 3(4), 101900. 10.1016/j.xpro.2022.101900

35. Ranjan, A., Nguyen, V. Q., Liu, S., Wisniewski, J., Kim, J. M., Tang, X., Mizuguchi, G., Elalaoui, E., Nickels, T. J., Jou, V., English, B. P., Zheng, Q., Luk, E., Lavis, L. D., Lionnet, T., & Wu, C. (2020). Live-cell single particle imaging reveals the role of RNA polymerase II in histone H2A.Z eviction. eLife, 9, e55667. 10.7554/eLife.55667

36. Saravanan, B., Soota, D., Islam, Z., Majumdar, S., Mann, R., Meel, S., Farooq, U., Walavalkar, K., Gayen, S., Singh, A. K., Hannenhalli, S., & Notani, D. (2020). Ligand dependent gene regulation by transient ERα clustered enhancers. PLOS Genetics, 16(1), e1008516. 10.1371/journal.pgen.1008516

37. Sigova, A. A., Abraham, B. J., Ji, X., Molinie, B., Hannett, N. M., Guo, Y. E., Jangi, M., Giallourakis, C. C., Sharp, P. A., & Young, R. A. (2015). Transcription factor trapping by RNA in gene regulatory elements. Science, 350(6263), 978–981. 10.1126/science.aad3346

38. Soota, D., Saravanan, B., Mann, R., Kharbanda, T., & Notani, D. (2024). RNA fine-tunes estrogen receptor-alpha binding on low-affinity DNA motifs for transcriptional regulation. The EMBO Journal, 1–25. 10.1038/s44318-024-00225-y

39. Soutourina, J., Bordas-Le Floch, V., Gendrel, G., Flores, A., Ducrot, C., Dumay-Odelot, H., Soularue, P., Navarro, F., Cairns, B. R., Lefebvre, O., & Werner, M. (2006). Rsc4 Connects the Chromatin Remodeler RSC to RNA Polymerases. Molecular and Cellular Biology, 26(13), 4920–4933. 10.1128/MCB.00415-06

40. Stenoien, D. L., Mancini, M. G., Patel, K., Allegretto, E. A., Smith, C. L., & Mancini, M. A. (2000). Subnuclear trafficking of estrogen receptor-alpha and steroid receptor coactivator-1. *Molecular Endocrinology (Baltimore*, Md.), 14(4), 518–534. 10.1210/mend.14.4.0436

41. Swinstead, E. E., Miranda, T. B., Paakinaho, V., Baek, S., Goldstein, I., Hawkins, M., Karpova, T. S., Ball, D., Mazza, D., Lavis, L. D., Grimm, J. B., Morisaki, T., Grøntved, L., Presman, D. M., & Hager, G. L. (2016). Steroid Receptors Reprogram FoxA1 Occupancy through Dynamic Chromatin Transitions. Cell, 165(3), 593–605. 10.1016/j.cell.2016.02.067

42. Veloso, A., Kirkconnell, K. S., Magnuson, B., Biewen, B., Paulsen, M. T., Wilson, T. E., & Ljungman, M. (2014). Rate of elongation by RNA polymerase II is associated with specific gene features and epigenetic modifications. Genome Research, 24(6), 896–905. 10.1101/gr.171405.113

43. Vos, S. M., Farnung, L., Urlaub, H., & Cramer, P. (2018). Structure of paused transcription complex Pol II–DSIF–NELF. Nature, 560(7720), 601–606. 10.1038/s41586-018-0442-2

44. Welboren, W.-J., van Driel, M. A., Janssen-Megens, E. M., van Heeringen, S. J., Sweep, F. C., Span, P. N., & Stunnenberg, H. G. (2009). ChIP-Seq of ERalpha and RNA polymerase II defines genes differentially responding to ligands. The EMBO Journal, 28(10), 1418–1428. 10.1038/emboj.2009.88

45. Xu, Y., Huangyang, P., Wang, Y., Xue, L., Devericks, E., Nguyen, H. G., Yu, X., Oses-Prieto, J. A., Burlingame, A. L., Miglani, S., Goodarzi, H., & Ruggero, D. (2021). ERα is an RNA-binding protein sustaining tumor cell survival and drug resistance. Cell, 184(20), 5215–5229.e17. 10.1016/j.cell.2021.08.036

46. Yague-Sanz, C., Migeot, V., Larochelle, M., Bachand, F., Wéry, M., Morillon, A., & Hermand, D. (2023). Chromatin remodeling by Pol II primes efficient Pol III transcription. Nature Communications, 14(1), 3587. 10.1038/s41467-023-39387-4

47. Zeitlinger, J., Stark, A., Kellis, M., Hong, J.-W., Nechaev, S., Adelman, K., Levine, M., & Young, R. A. (2007). RNA polymerase pausing at developmental control genes in the Drosophila melanogaster embryo. Nature Genetics, 39(12), 1512–1516. 10.1038/ng.2007.26

48. Zhang, Y., Liu, T., Meyer, C. A., Eeckhoute, J., Johnson, D. S., Bernstein, B. E., Nusbaum, C., Myers, R. M., Brown, M., Li, W., & Liu, X. S. (2008). Model-based Analysis of ChIP-Seq (MACS). Genome Biology, 9(9), R137. 10.1186/gb-2008-9-9-r137

