## Supplementary material for "Paused Polymerase synergises with transcription factors and RNA to enhance the transcriptional potential of promoters": Supp Figures

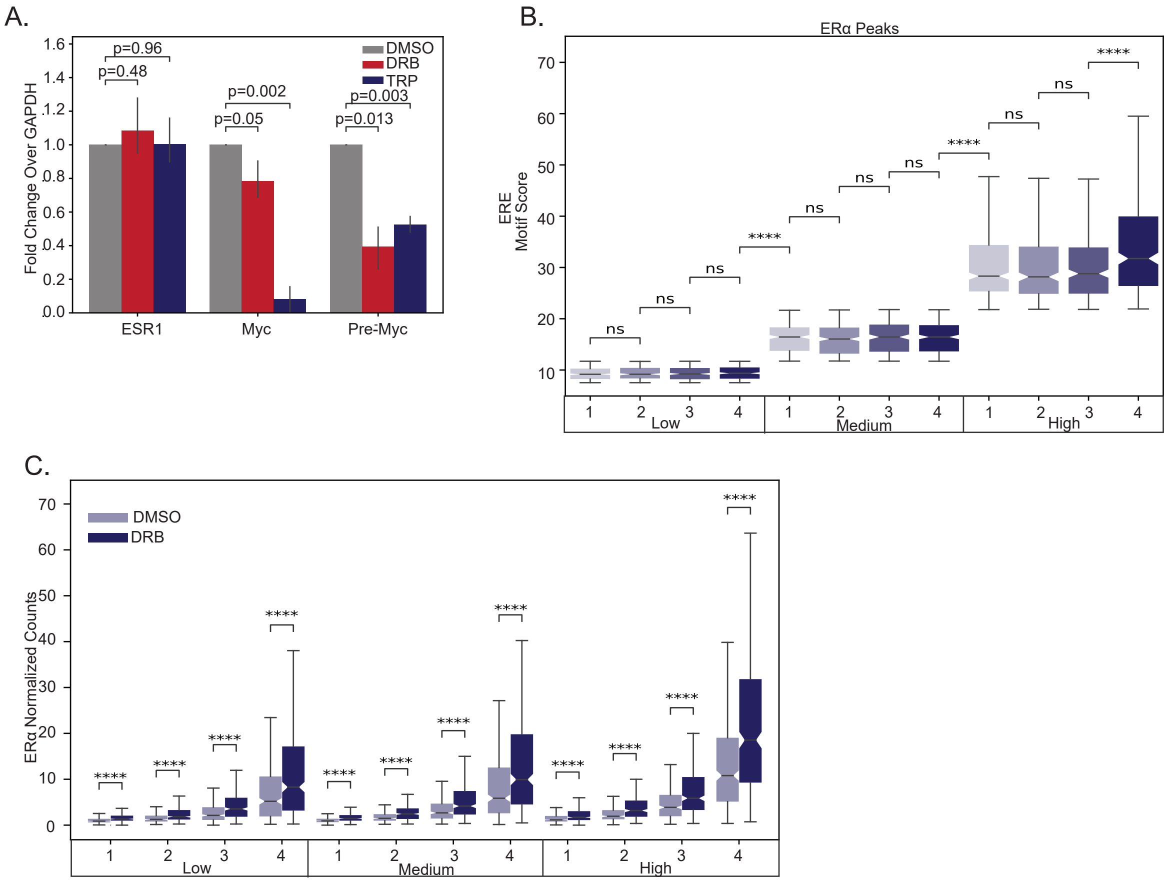
A.
Fold Change Over GAPDH
DMSO
DRB
TRP
p=0.96
p=0.48
p=0.002
p=0.05
p=0.003
p=0.013
ESR1
Myc
Pre-Myc
B.
ERα Peaks
ERE Motif Score
ns
****
Low
Medium
High
C.
DMSO
DRB
ERα Normalized Counts
****
Low
Medium
High

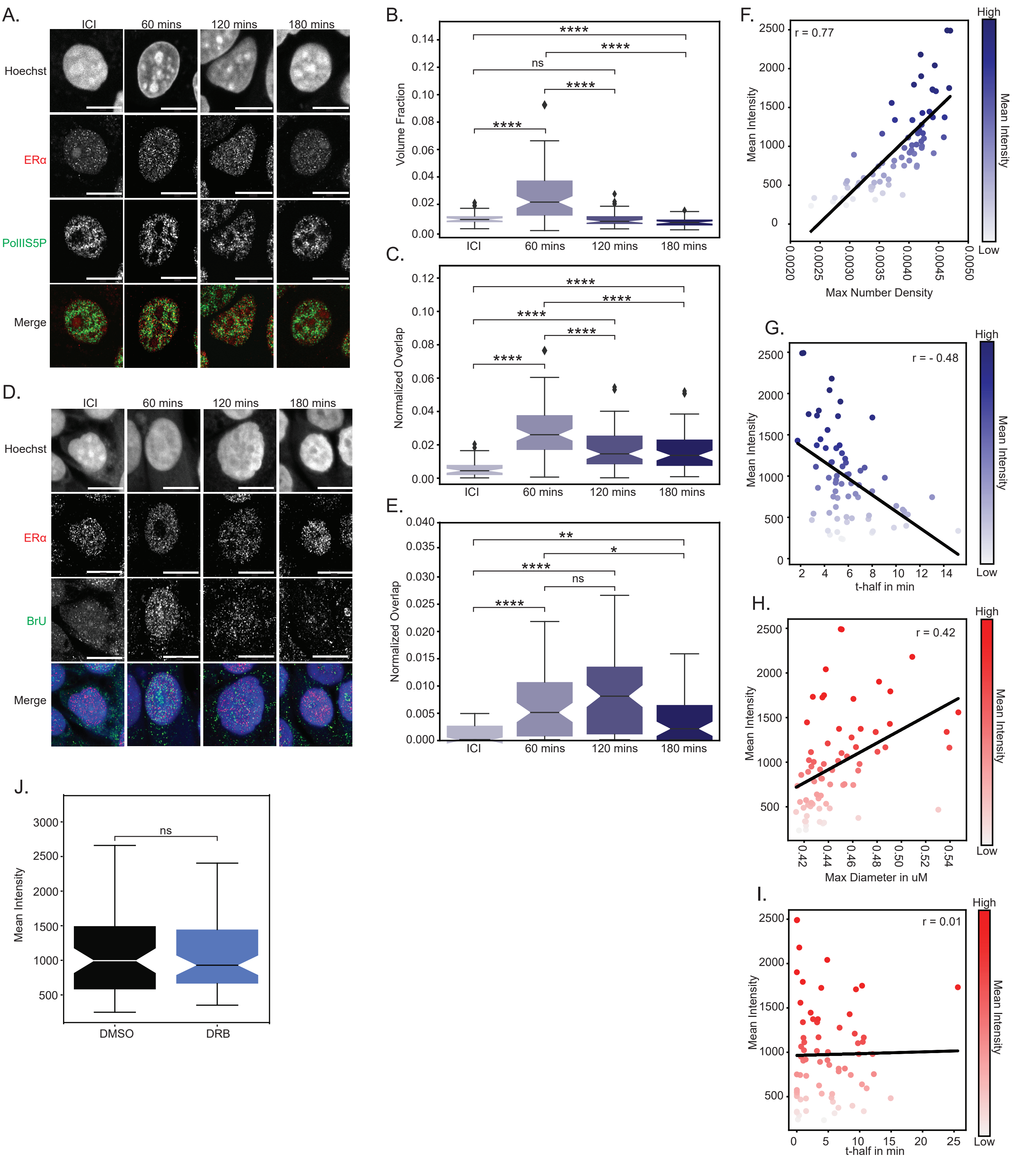
A.
ICI
60 mins
120 mins
180 mins
Hoechst
ERα
PolIIS5P
Merge
B.
Volume Fraction
ns
ICI
60 mins
120 mins
180 mins
C.
Normalized Overlap
D.
BrU
E.
F.
r = 0.77
Mean Intensity
Max Number Density
High
Low
G.
r = - 0.48
t-half in min
H.
r = 0.42
Max Diameter in uM
I.
r = 0.01
J.
DMSO
DRB

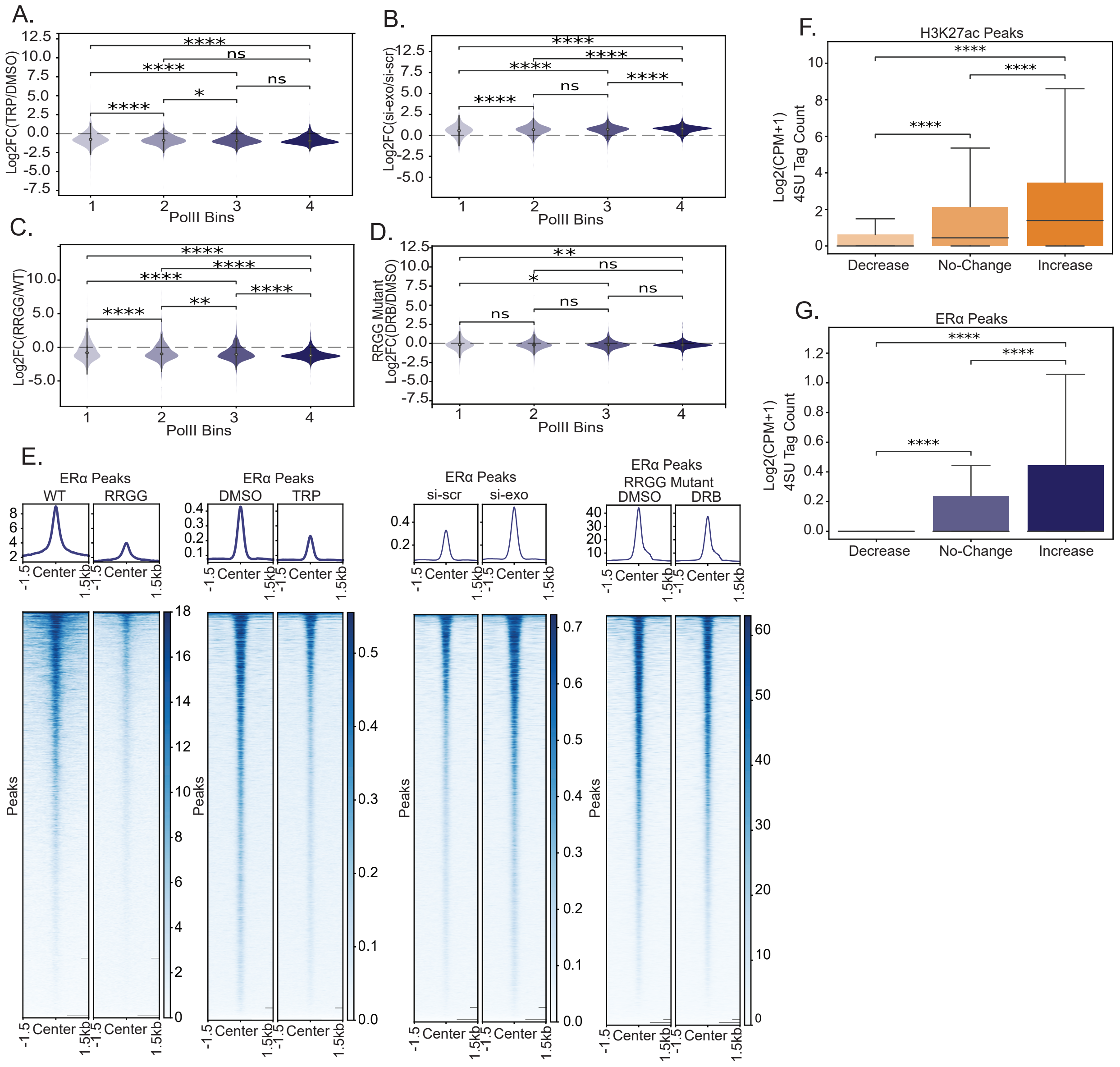

A.
Log2FC(TRP/DMSO)
PolII Bins
B.
Log2FC(si-exo/si-scr)
PolII Bins
C.
Log2FC(RRGG/WT)
PolII Bins
D.
RRGG Mutant
Log2FC(DRB/DMSO)
PolII Bins
E.
ERα Peaks
WT
RRGG
ERα Peaks
DMSO
TRP
ERα Peaks
si-scr
si-exo
ERα Peaks
RRGG Mutant
DMSO
DRB
Peaks
Center
F.
H3K27ac Peaks
Log2(CPM+1)
4SU Tag Count
Decrease
No-Change
Increase
G.
ERα Peaks
Log2(CPM+1)
4SU Tag Count
Decrease
No-Change
Increase

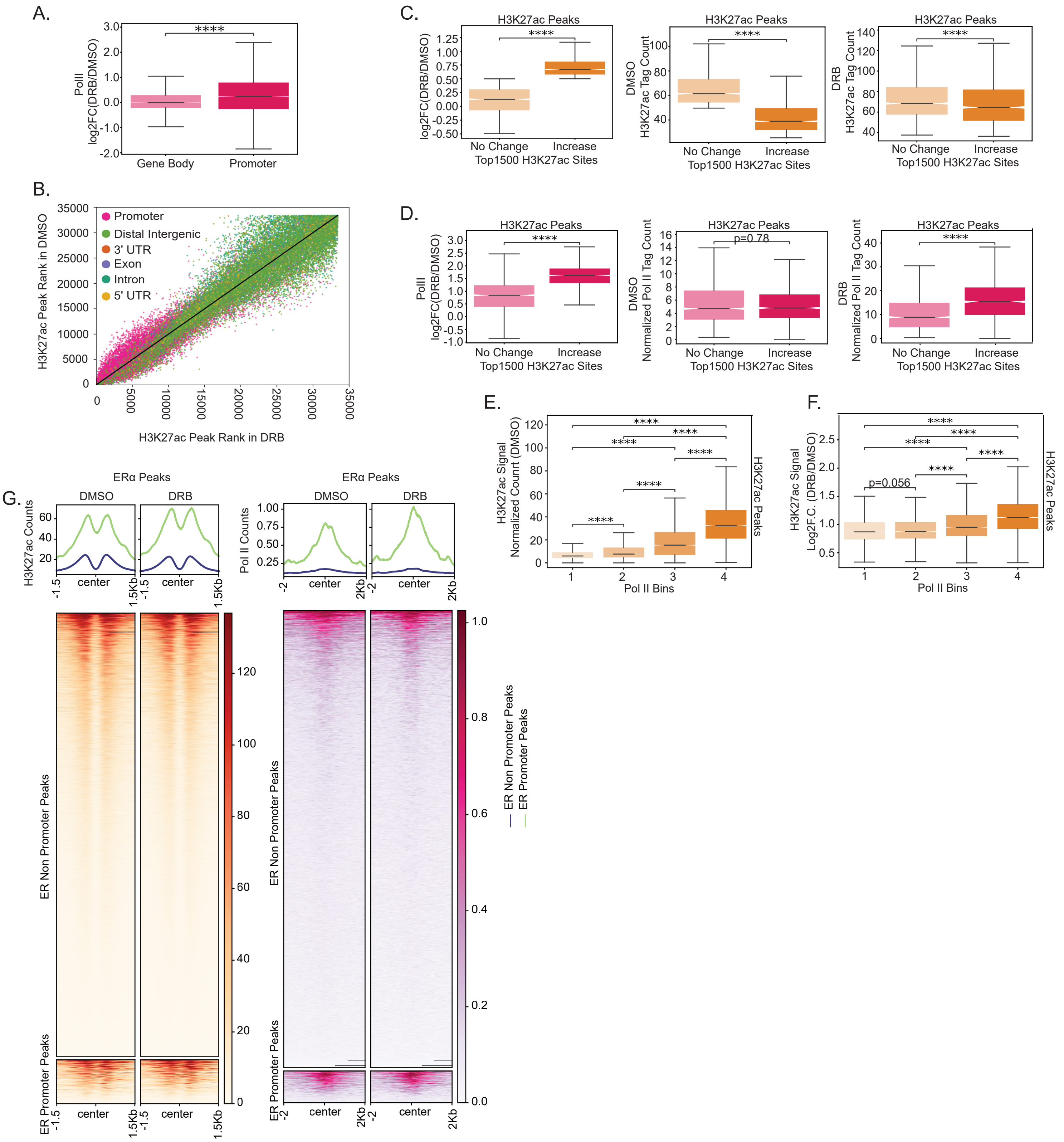

A.
PolII log2FC(DRB/DMSO)
****
Gene Body
Promoter
B.
H3K27ac Peak Rank in DMSO
H3K27ac Peak Rank in DRB
Promoter
Distal Intergenic
3' UTR
Exon
Intron
5' UTR
C.
H3K27ac Peaks
log2FC(DRB/DMSO)
DMSO H3K27ac Tag Count
DRB H3K27ac Tag Count
No Change
Increase
Top1500 H3K27ac Sites
D.
H3K27ac Peaks
PolII log2FC(DRB/DMSO)
DMSO Normalized Pol II Tag Count
DRB Normalized Pol II Tag Count
p=0.78
No Change
Increase
Top1500 H3K27ac Sites
E.
H3K27ac Signal Normalized Countt (DMSO)
H3K27ac Peaks
Pol II Bins
F.
H3K27ac Signal Log2F.C. (DRB/DMSO)
p=0.056
H3K27ac Peaks
Pol II Bins
G.
ERα Peaks
DMSO
DRB
H3K27ac Counts
Pol II Counts
center
ER Non Promoter Peaks
ER Promoter Peaks

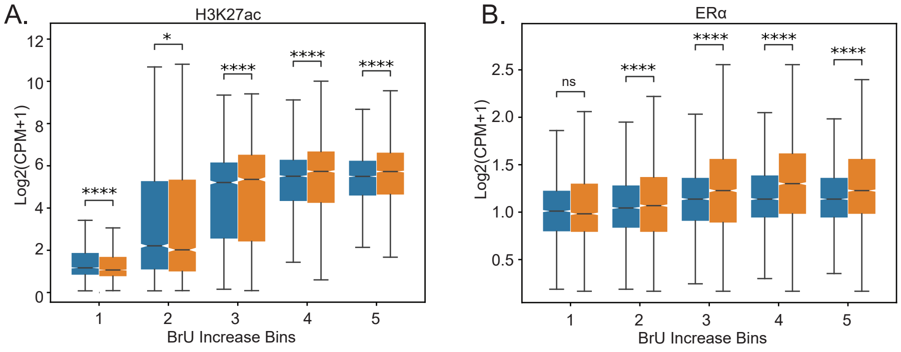
A.
H3K27ac
Log2(CPM+1)
12
10
8
6
4
2
0
****
*
****
****
****
1
2
3
4
5
BrU Increase Bins
B.
ERα
Log2(CPM+1)
2.5
2.0
1.5
1.0
0.5
ns
****
****
****
****
1
2
3
4
5
BrU Increase Bins

**Table 1: Oligos for ERα-Halo Cloning**

| Name | Sequence |
|---|---|
| EFS_Fragment_FW | ggtctcTCTAGAtgatgcgggcccttcgagtggctc |
| EFS_Fragment_RW | ggtctcttccatggtggcgccagcctgcacgcgttcacc |
| Halo_Gene_Fragment1_FW | GGTCTCATGGAAATCGGTACTGGCTTTCCATTCGACCCC |
| Halo_Gene_Fragment1_RW | ggtctcGGTTTCGCGGGCAAATTCTGGCCATT |
| Halo_Gene_Fragment2_FW | GGTCTCGAAACCTTCCAGGCCTTCCGCACCACC |
| Halo_Gene_Fragment2_RW | GGTCTCACCGGAAATCTCCAGAGTAGACAGCCAGCG |
| ER_Gene_Fragement1_FW | GGTCTCACCGGATCCGGACTCAGATCTCGAGCTCAAGCTTCG |
| ER_Gene_Fragement1_FW | ggtctcGGGCTCGTTCTCCAGGTAGTAGGGC |
| ER_Gene_Fragement2_FW | ggtctcGAGCCCAGCGGCTACACGGTGCGCGaggc |
| ER_Gene_Fragement2_FW | ggtctcGCAGCTCTTCCTCCTGTTTTTATCAATGGT |
| ER_Gene_Fragement3_FW | GGTCTCAGCTGCCAGGCCTGCCGGCTCCGCAAA |
| ER_Gene_Fragement3_FW | GGTCTCTAGACCAATCATCAGGATCTCTAGCC |
| ER_Gene_Fragement4_FW | GGTCTCGGTCTAGTCTGGCGCTCCATGGAGCA |
| ER_Gene_Fragement4_FW | GGTCTCCAGGTCATAGAGGGGCACCACGTTC |
| ER_Gene_Fragement5_FW | GGTCTCGACCTGCTGCTGGAGATGCTGGACG |
| ER_Gene_Fragement5_FW | GGTCTCCGGCCGTAAGATACATTGATGAGTTTGGACAaa |

**Table 2: Oligos for qPCR**

| Name | Sequence |
|---|---|
| ESR1_qPCR_FW | TGTGTCCAGCCACCA ACC AG |
| ESR1_qPCR_RW | TTCAAC ATTCTCCCTCCTCTTCGG |
| Myc_qPCR_FW | GAGGAGACATGGTGAACCAGAG |
| Myc_qPCR_RW | CCAGCTTCTCTGAGACGAGC |
| PreMyc_qPCR_FW | CTTCTCTCCGTCCTCGGATTC |
| PreMyc_qPCR_RW | CCTTCCTAATAAGAGTGGCCCG |
| GAPDH_FW | CGCTCTCTGCTCCTCCTGTT |
| GAPDH_RW | CCATGGTGTCTGAGCGATGT |

### Table 3: NGS Data with Accession Number

| Accession Number | Dataset | Source |
|---|---|---|
| GSM678539 | GRO-Seq-40mins-E2 | (Hah et al., 2011) |
| GSM365930 | Pol II_ChiP_seq_MCF7_1hr_E2 | (Welboren et al., 2009) |
| GSM1338772 | DRB_BrU_seq_Mock | (Veloso et al., 2014) |
| GSM1338773 | DRB_BrU_seq_DRB_Release | (Veloso et al., 2014) |
| PRJNA691029 | 4SU_Seq_DRB_MCF7 | (Beckman et al., 2021) |
| GSM855XXXX | DMSO_control_DRB_ERalpha_Rep1 | This Study |
| GSM855XXXX | DMSO_control_DRB_ERalpha_Rep2 | This Study |
| GSM855XXXX | DMSO_control_DRB_Flag_RRGG_ER_Rep1 | This Study |
| GSM855XXXX | DMSO_control_DRB_Flag_RRGG_ER_Rep2 | This Study |
| GSM855XXXX | DMSO_control_DRB_H3K27ac_Rep1 | This Study |
| GSM855XXXX | DMSO_control_DRB_H3K27ac_Rep2 | This Study |
| GSM855XXXX | DMSO_control_DRB_Pol II | This Study |
| GSM855XXXX | DMSO_control_TRP_ERalpha_Rep1 | This Study |
| GSM855XXXX | DMSO_control_TRP_ERalpha_Rep2 | This Study |
| GSM855XXXX | DRB_treatment_DRB_ERalpha_Rep1 | This Study |
| GSM855XXXX | DRB_treatment_DRB_ERalpha_Rep2 | This Study |
| GSM855XXXX | DRB_treatment_DRB_Flag_RRGG_ER_Rep1 | This Study |
| GSM855XXXX | DRB_treatment_DRB_Flag_RRGG_ER_Rep2 | This Study |
| GSM855XXXX | DRB_treatment_DRB_H3K27ac_Rep1 | This Study |
| GSM855XXXX | DRB_treatment_DRB_H3K27ac_Rep2 | This Study |
| GSM855XXXX | DRB_treatment_DRB_Pol II | This Study |
| GSM855XXXX | TRP_treatment_TRP_ERalpha_Rep1 | This Study |
| GSM855XXXX | TRP_treatment_TRP_ERalpha_Rep2 | This Study |
| GSM855XXXX | si_scr_ERalpha | This Study |
| GSM855XXXX | si_exo_ERalpha | This Study |
